# The *Arabidopsis thaliana* PeptideAtlas; harnessing world-wide proteomics data for a comprehensive community proteomics resource

**DOI:** 10.1101/2021.05.03.442425

**Authors:** Klaas J. van Wijk, Tami Leppert, Qi Sun, Sascha S. Boguraev, Zhi Sun, Luis Mendoza, Eric W. Deutsch

## Abstract

We developed a new resource, the Arabidopsis PeptideAtlas (www.peptideatlas.org/builds/arabidopsis/), to solve central questions about the Arabidopsis proteome, such as the significance of protein splice forms, post-translational modifications (PTMs), or simply obtain reliable information about specific proteins. PeptideAtlas is based on published mass spectrometry (MS) analyses collected through ProteomeXchange and reanalyzed through a uniform processing and metadata annotation pipeline. All matched MS-derived peptide data are linked to spectral, technical and biological metadata. Nearly 40 million out of ∼143 million MSMS spectra were matched to the reference genome Araport11, identifying ∼0.5 million unique peptides and 17858 uniquely identified proteins (only isoform per gene) at the highest confidence level (FDR 0.0004; 2 non-nested peptides ≥ 9 aa each), assigned canonical proteins, and 3543 lower confidence proteins. Physicochemical protein properties were evaluated for targeted identification of unobserved proteins. Additional proteins and isoforms currently not in Araport11 were identified, generated from pseudogenes, alternative start, stops and/or splice variants and sORFs; these features should be considered for updates to the Arabidopsis genome. Phosphorylation can be inspected through a sophisticated PTM viewer. This new PeptideAtlas is integrated with community resources including TAIR, tracks in JBrowse, PPDB and UniProtKB. Subsequent PeptideAtlas builds will incorporate millions more MS data.

**One sentence summary:** A new web resource providing the global community with mass spectrometry-based Arabidopsis proteome information and its spectral, technical and biological metadata integrated with TAIR and JBrowse

## INTRODUCTION

*Arabidopsis thaliana* (from here on Arabidopsis) was the first plant species for which the nuclear genome was sequenced and it has served as a model species for plant biology research for the last ∼25 years (Provart et al., 2016). The current Arabidopsis genome release version 11 (Araport11) contains 27,655 protein-coding gene loci represented by 48,359 transcripts (Cheng et al., 2017). The collective set of proteins in Arabidopsis, referred to as the proteome, carries out essential functions in metabolism, gene expression, signal transduction, transport and more. The proteome not only varies with time, development, and (a)biotic conditions, but also undergoes a wide range of dynamic reversible and irreversible post-translational modifications (PTMs; *e.g*. phosphorylation, ubiquitination, acetylation). Furthermore, proteins are distributed across subcellular locations, such as the various organelles, and many proteins often stably or dynamically interact with other proteins. Whereas genome sequencing technologies combined with large scale RNAseq data and computation can predict the theoretical set of protein coding genes in an organism, cell-type specific and subcellular protein abundance, protein PTMs, and protein interactions cannot be predicted but must be experimentally determined at the protein level. Furthermore, even the best annotated genomes such as Arabidopsis and human cannot easily predict which mRNA splice forms result in proteins; indeed the impact of alternative splicing on the human (and other species) proteome is still under debate (Blencowe, 2017; Tress et al., 2017). The use of proteomics data for plant genome annotation has only very recently begun to make a more systematic impact under the term ‘proteogenomics’ (Castellana et al., 2014; Walley and Briggs, 2015; Chapman and Bellgard, 2017; Zhu et al., 2017; Ren et al., 2019). This has included genomes of Arabidopsis (Zhu et al., 2017; Zhang et al., 2019), rice (Ren et al., 2019; Chen et al., 2020), maize (Castellana et al., 2014), grape (Chapman and Bellgard, 2017) and sweet potato (Al-Mohanna et al., 2019).

Initial MS-based plant proteomics studies appeared in the year 2000, investigating proteomes of maize and pea (Chang et al., 2000; Peltier et al., 2000; van Wijk, 2000) at a time when there were no sequenced plant genomes, instead relying on expressed sequence tag (EST) assemblies. With the release of the first partial Arabidopsis (ecotype Columbia-0) genome sequence (Initiative, 2000), Arabidopsis rapidly became the organism of choice for plant proteomics studies. Initially, MS was used for the study of subcellular organelles such as chloroplasts and mitochondria (Millar et al., 2001; Peltier et al., 2002; Schubert et al., 2002; Ytterberg, 2002), plant structures such as pollen (Mayfield et al., 2001), and protein complexes (Peltier et al., 2001). MS-based proteomics has since become increasingly successful for studying proteomes of different plant organs, cell types, and (subcellular) compartments as well as the many plant protein PTMs, such as phosphorylation (Stecker et al., 2014; Balmant et al., 2016), lysine acetylation (Hosp et al., 2017), ubiquitination (Vierstra, 2012) and SUMOylation (Miura and Hasegawa, 2010), redox modifications (Akter et al., 2015; Waszczak et al., 2015), N-terminal acetylation (Rowland et al., 2015; Willems et al., 2017) or lysine acetylation (Hartl et al., 2017). For recent reviews on PTMs in plants, see (Friso and van Wijk, 2015; Millar et al., 2019). Proteomics has also been extensively used to study plant responses to (a)biotic conditions, and plant developmental processes in *e.g*. roots, seeds, and leaves, reviewed in (Vanderschuren et al., 2013; Ruiz-May et al., 2019). The progress of proteomics research of plants has been regularly reviewed, mostly in an attempt to consolidate plant proteome information, including protein detection, various PTMs, abundance measurements, and to provide updates of plant proteomics and mass spectrometry technologies and plant proteome databases (Tan et al., 2017; Misra, 2018). A range of plant proteome databases by individual labs have been developed, mostly for Arabidopsis proteins, typically focused quite narrowly towards a particular aspect of plant proteomics, such as subcellular compartments (San Clemente and Jamet, 2015; Salvi et al., 2018), protein location (SUBA and PPDB) (Sun et al., 2009; Tanz et al., 2013), or PTMs (Schulze et al., 2015; Willems et al., 2019). Most recently, a more comprehensive Arabidopsis proteome database (ATHENA) was released to mine a large scale experimental proteome data set involving multiple tissue types (Mergner et al., 2020).

The global scientific community has developed a wide range of initiatives to capture and store highly data-rich MS-based proteomics information using standardized bioinformatics workflows and file formats (Orchard et al., 2003; Deutsch et al., 2017a). The ProteomeXchange consortium (http://www.proteomexchange.org/) coordinates standard data submission and dissemination pipelines across the main proteomics repositories and promotes submission of all published datasets and open data policies in the field (Vizcaino et al., 2014; Deutsch et al., 2017b; Deutsch et al., 2020). The consortium has made tremendous progress in getting the community to deposit its datasets in conjunction with publication of an article. Currently there are well over 15000 released ProteomeXchange datasets (PXDs). Many plant journals such as The Plant Cell, Plant Physiology, Plant Journal, Molecular Plant and others, strongly encourage MS data deposition for publications that rely on MS-based proteomics. Currently (at the time of submission), ProteomeXchange has over 1200 released PXDs for proteome datasets from many plant species (and a few from algae) of which ∼425 PXDs are from *Arabidopsis*.

PeptideAtlas (http://www.peptideatlas.org/) reprocesses MS datasets available through ProteomeXchange with the Trans-Proteomic Pipeline (TPP) (Keller et al., 2005; Deutsch et al., 2015; Slagel et al., 2015), and makes an integrated view of the results available to the community. So far, PeptideAtlas has focused heavily on the human proteome with a first publication in 2005 (Desiere et al., 2005) and ongoing contributions to the Human Proteome Project (HPP) including yearly advances in coverage of the human proteome (Omenn et al., 2019; Omenn et al., 2020). However, PeptideAtlas has also created builds for several other species, including pig (*Sus scrofa*) (Hesselager et al., 2016), chicken (*Gallus gallus*) (McCord et al., 2017), cow (*Bos Taurus*) (Bislev et al., 2012), the pathogens *Pseudomonas aeruginosa* (33757883) and *Candida albicans* (Vialas et al., 2014) and the yeast *Saccharomyces cerevisiae* (King et al., 2006). Yet there have not previously been PeptideAtlas builds for any plant species. Given the significant amount of PXD submissions for plants, and in particular Arabidopsis, this provides a unique opportunity to take full advantage of the rapidly growing amounts of MS-based proteomics data for Arabidopsis, to build a thorough understanding of the observed Arabidopsis proteome.

The current study is the first report on a project that will take advantage of the current and anticipated submissions to ProteomeXchange by reanalyzing these data through the TPP to generate PeptideAtlas builds for Arabidopsis and in later stages additional plant species. This freely available Arabidopsis PeptideAtlas provides the global community with high quality, fully reprocessed MS-based proteome information together with its metadata. This resource can be used to solve central questions about the Arabidopsis proteome, such as the significance of protein splice forms, PTMs, or simply obtain reliable information about specific protein sets of interest, without the need to be an expert in MS. The Arabidopsis PeptideAtlas provides immediate insight into: i) which Arabidopsis proteins have been identified and with how much protein sequence coverage, ii) relative protein abundance based on frequency of observations across datasets and sampling across plant organs, cell types, organelles, (a)biotic treatments, development and complexes, iii) enrichment for specific post-translational modifications, iv) which proteins have not yet been observed (the ‘dark’ proteome) and v) specific information to improve genome annotation, including discovery of protein-coding small ORFs. We envision that these Arabidopsis PeptideAtlas builds will stimulate labs around the world to submit their proteomics and MS data to ProteomeXchange, further accelerating our knowledge about the expression and PTMs of plant proteins.

## METHODS

### Selection and downloads of ProteomeXchange submissions

PXDs were selected based on several criteria, including mass spectrometer type with preference for Orbitrap-type instruments from Thermo (Q Exactive models, LTQ-Orbitrap Velos/Elite, Orbitrap Fusion Lumos), submissions from 2018 and 2019, and samples including subcellular fractions or specific PTMs. The rationale is provided in the RESULTS & DISCUSSION section. Raw files for the selected PXDs were downloaded from ProteomeXchange. Supplemental Dataset 1 provides the final 52 selected PXDs and information about instrument, sample (*e.g.* subcellular proteome, plant organ), number of raw files and MSMS spectra (searched and matched), identified proteins and peptides, submitting lab and associated publication, as well as several informative key words.

### Extraction and annotation of metadata

For each selected dataset, we obtained information associated with the submission, as well as the publication if available. This information was used to determine search parameters and provide meaningful tags that describe the samples in some detail. These tags are visible for the relevant proteins in the PeptideAtlas. If needed, we contacted the submitters for more information about the raw files. To facilitate the metadata assignments and association to specific raw files, we developed a metadata annotation system that is aimed to provide detailed information to each matched spectrum for the users of the PeptideAtlas. Where possible, we incorporated controlled vocabularies for plant parts and developmental stages, growth conditions, sample purification methods, as well as protein/peptide labeling and processing steps (*e.g.* type of enzyme used for generation of peptides). These controlled vocabularies are from the Planteome (PO, PECO) (https://github.com/Planteome), Gene Ontology (http://geneontology.org/), as well as PSI-MS (http://www.psidev.info/groups/mass-spectrometry) (Mayer et al., 2013), Unimod (https://www.unimod.org) (Creasy and Cottrell, 2004), PSI-MOD (https://www.ebi.ac.uk/ols/ontologies/mod) (Montecchi-Palazzi et al., 2008), and the Experimental Factor Ontology (EFO) (https://www.ebi.ac.uk/ols/ontologies/efo).

### Assembly of protein search space

We assembled a comprehensive protein search space comprising the predicted *Arabidopsis* protein sequences from i) Araport11 (Cheng et al., 2017), ii) TAIR10 (Lamesch et al., 2012), iii) UniProtKB (UniProt, 2020), iv) RefSeq (https://www.ncbi.nlm.nih.gov/refseq) (Li et al., 2020), v) from the repository ARA-PEPs (http://www.biw.kuleuven.be/CSB/ARA-PEPs) (Hazarika et al., 2017) with 13748 putative peptides (small Open Reading Frames (sORFs) and low molecular weight proteins (LWs)) as well as 341 novel stress-induced peptides (SIPs) currently not annotated in TAIR10 or Araport11, vi) from Dr Eve Wurtele (Iowa State university) assembled based on RNAseq data, vii) GFP, RFP and YFP protein sequences commonly used as reporters and affinity enrichments, viii) 116 contaminant protein sequences frequently observed in proteome samples (*e.g.* keratins, trypsin, BSA) (https://www.thegpm.org/crap/). Table 1 shows the number of sequences for each set, their overlap and unique protein sequences.

**Table 1.**
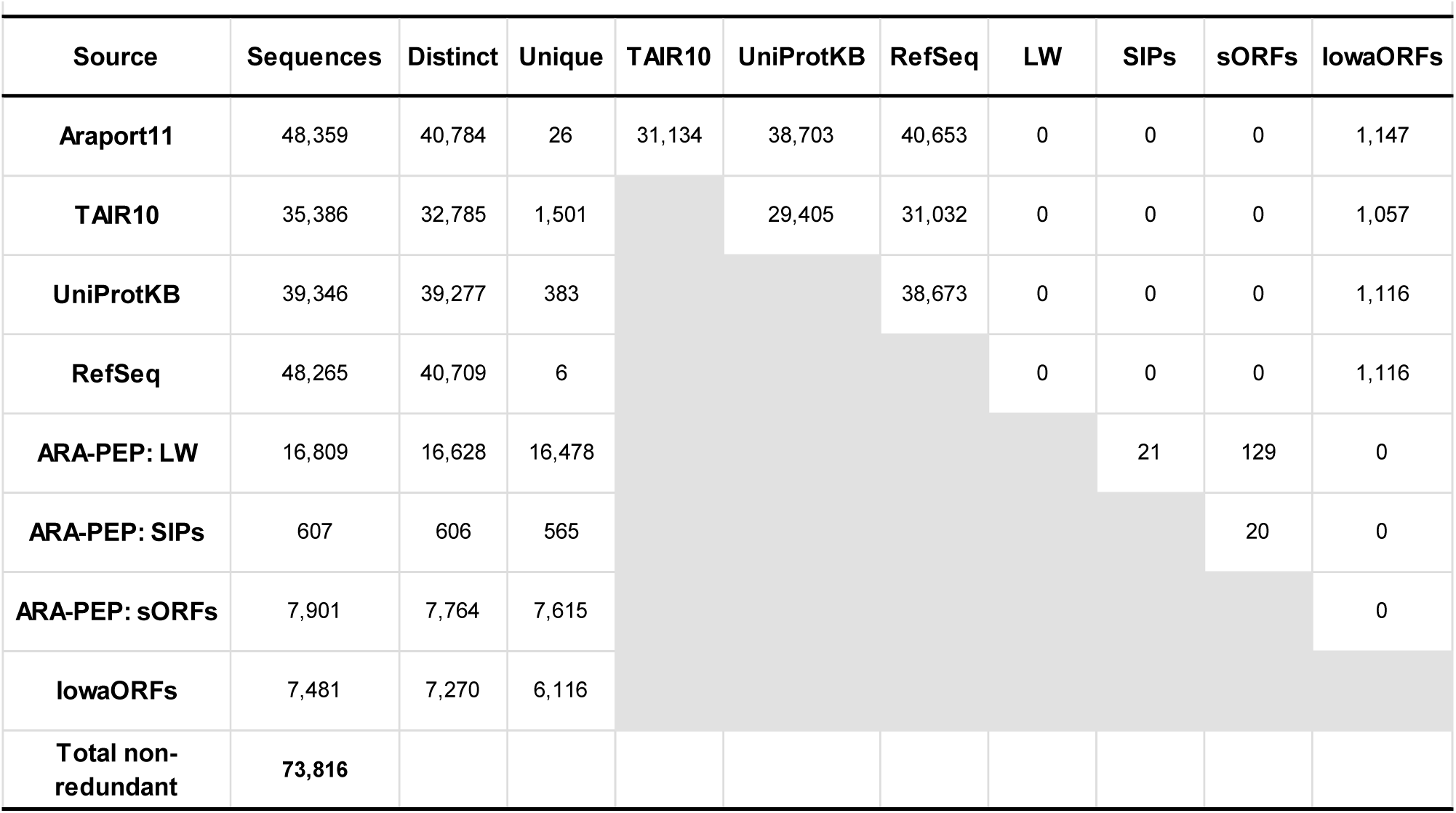
The assembly of protein sequences from different sources used as the protein search space, and the respective number of total, distinct, and unique sequences in each source, as well as the sequence-identical intersection among sources.

### The TPP data processing pipeline

For all selected datasets, the vendor-format raw files were downloaded from the hosting ProteomeXchange repository, converted to mzML files (Martens et al., 2011) using ThermoRawFileParser (Hulstaert et al., 2020) for Thermo Fisher Scientific instruments or the msconvert tool for SCIEX wiff files from the ProteoWizard toolkit (Chambers et al., 2012), and then analyzed with the TPP. The TPP analysis consisted of sequence database searching with Comet (Eng and Deutsch, 2020) and post-search validation with several additional TPP tools as follows: PeptideProphet (Keller et al., 2002) was run to assign probabilities of being correct for each peptide-spectrum match (PSM) using semi-parametric modeling of the Comet expect scores with z-score accurate mass modeling of precursor m/z deltas. These probabilities were further refined via corroboration with other PSMs, such as multiple PSMs to the same peptide sequence but different peptidoforms or charge states, using the iProphet tool (Shteynberg et al., 2011).

For datasets in which trypsin was used as the protease to cleave proteins into peptides, two parallel searches were performed, one with full tryptic specificity and one with semi-tryptic specificity. The semi-tryptic searches were carried out with the following possible variable modifications (5 max per peptide): oxidation of Met or Trp (+15.9949), acetylation of Lys (+42.0106), peptide N-terminal Gln to pyro-Glu (−17.0265), peptide N-terminal Glu to pyro-Glu (−18.0106), deamidation of Asn or Gln (+0.9840), peptide N-term acetylation (+42.0106), and if peptides were specifically affinity enriched for phosphopeptides, also phosphorylation of Ser, Thr or Tyr (+79.9763). For the full tryptic searches, we also added oxidation of Pro or His, formylation of peptide N-termini, Ser, or Thr (+27.9949). In both searches, fixed modifications for carbamidomethylation of Cys (+57.0215) if treated with reductant and iodoacetamide and isobaric tag modifications (TMT, iTRAQ) were applied as appropriate. Both variable and fixed modifications were applied to dimethyl labeled datasets as appropriate. Four missed cleavages were allowed (RP or KP do not count as a missed cleavage). Several datasets were generated using other proteases (GluC, ArgC, Chymotrypsin); these data sets were processed similarly to those generated by trypsin with the exception that the relevant enzyme was chosen. Some of the datasets contain the analysis of extracted peptidomes in which no protease treatment was used and these datasets were searched with ‘no enzyme’.

### PeptideAtlas Assembly

In order to create the combined PeptideAtlas build of all experiments, all datasets were thresholded at a probability that yields an iProphet model-based FDR of 0.001 at the peptide level. The exact probability varies from experiment to experiment depending on how well the modeling can separate correct from incorrect. This probability threshold is typically greater than 0.99. As more and more experiments are combined, the total FDR increases unless the threshold is made more stringent (Deutsch et al., 2016b). The final iProphet model-based peptide sequence level FDR across all experiments is 0.001, corresponding to a PSM-level False Discover Rate (FDR) of 0.0001. Throughout the procedure, decoy identifications are retained and then used to compute final decoy-based FDRs. The decoy-based PSM-level FDR is 0.0001 (4843 decoy PSMs out of 40 million), peptide sequence-level FDR is 0.001 (746 decoy sequences out of 535,000), and the final protein-level FDR is 0.03 (683 decoy proteins out of 21297). This is a very stringent threshold, which has the unfortunate effect of discarding a high portion of correct identifications. This is unavoidable if the final build is to maintain a low number of false positives, because while many correct identifications are discarded, they are mixed indistinguishably with false positives at a rate higher than is desirable.

### Protein identification confidence levels and classification

Proteins are identified at different confidence levels using standardized assignments to different confidence levels based on various attributes and relationships to other proteins using a relatively complex but precise ten-tier system developed over many years for the human proteome PeptideAtlas (Farrah et al., 2011) (Table 2A). We simplified this ten-tier system to a simpler four category system in (Table 2B), which is more accessible to non-experts, and use this to summarize most of our findings. For all protein identifications and categorizations, all peptides must first meet the stringent PSM threshold already described above. For both systems, the highest confidence level category is the “canonical” category (Tier 1), which requires at least two uniquely-mapping non-nested (one not inside the other) peptides of at least nine amino acids in length with a total coverage of at least 18 amino acids, as required by the HPP guidelines (Deutsch et al., 2019) (Table 2A,B). The decoy-based canonical protein FDR is 0.0005 (only 8 decoys remain out of 18,045 canonical sequences including contaminants and contributed sequences).

**Table 2.** Protein identification confidence tiers and categories in the Arabidopsis PeptideAtlas build.

| Category | Definition |
| --- | --- |
| **Tier 1. Canonical** | Protein has at least two uniquely-mapping non-nested peptides of at least 9 residues with at least 18 residues of total coverage |
| **Tier 2. Indistinguishable representative** | Protein is selected as the representative of a set of proteins that are different in sequence but cannot be disambiguated based on the detected peptides. All peptides are shared with all group members. Others are "Indistinguishable". |
| **Tier 3. Representative** | Protein is selected a representative in a situation more complex than a set of indistinguishable, where several proteins have shared peptides and at least some of the proteins must have been detected but it is not possible to determine which ones. |
| **Tier 4. Marginally distinguished** | Protein that shares several peptides with a canonical protein, but also has one uniquely-mapping peptide that appears to distinguish it from the canonical. |
| **Tier 5. Weak** | Protein has at least one uniquely-mapping peptide of 9 residues in length but does not meet the criteria for canonical. |
| **Tier 6. Insufficient evidence** | Protein has one or more uniquely mapping peptides but none reach 9 residues in length. |
| **Tier 7. Indistinguishable** | Protein is part of a set of proteins that cannot be disambiguated and it not selected as a leader of the group. |
| **Tier 8. Subsumed** | Protein has only shared peptides and is not needed to explain the peptide evidence. |
| **Tier 9. Identical** | Protein has an identical protein sequence to another one, and this one is effectively removed from category competition. Its partner may be canonical. |
| **Tier 10. Not Observed** | Protein has no peptides above our PSM significance threshold. It may have low significant PSMs but there are not considered. |

| Category | Definition |
| --- | --- |
| **Canonical** (as in Tier 1 in Table 2A) | Protein has at least two uniquely-mapping non-nested peptides of at least 9 residues with at least 18 residues of total coverage |
| **Uncertain** (Tiers 2-7 in Table 2A) | Protein has too few uniquely-mapping peptides of $\geq 9$ aa to qualify for canonical status and may also have one or more shared peptides with other proteins. |
| **Redundant** (Tiers 8 & 9 in Table 2A) | Protein has only peptides that are can be assigned to other entries and thus these proteins are not needed to explain the observed peptide evidence. |
| **Not Observed** (Tier 10 in Table 2A) | Protein has no peptides above our PSM significance threshold. It may have low significance PSMs but they are not considered. |

#### The ten-tier system

When a group of proteins cannot be disambiguated because of shared peptides, one or more “leaders” of the group are designated as categories ‘indistinguishable representative’ (Tier 2) or ‘representative’ (Tier 3) (Table 2A). This means that the protein or one of its close siblings are detected, but it is not possible to disambiguate them. The “marginally distinguished” category (Tier 4) means that the protein shares peptides with a canonical entry but has some additional uniquely mapping peptide evidence that is however not sufficient to raise it to canonical level. The ‘weak’ category (Tier 5) (Table 2A) means that there is at least one uniquely mapping peptide that is nine or more residues long, but the evidence does not meet the criteria for being canonical. The ‘insufficient evidence’ category (Tier 6) means that all the uniquely mapping peptides are less than 9 residues long. While even one uniquely mapping peptide in theory uniquely identifies a protein, these guidelines guard against false positives due to our imperfect understanding of the reference proteome and incomplete b and y ion series identifications, which can lead to amino acid order transpositions and protein misassignment. Tiers 7, 8 and 9 describe proteins that share all their peptides with one or more proteins in an earlier tier, and thus are not needed to explain the available peptide evidence. Finally, all other proteins that lack any matched peptides observed above our minimum PSM significance threshold are categorized as ‘not observed’ proteins (Tier 10) (Table 2A).

#### The four-category system

In the simpler four-category system, proteins that have no uniquely mapping peptides but do not qualify as canonical (same as Tier 1) are categorized as ‘uncertain’ (Table 2B), corresponding to the sum of tiers 2-6 in Table 2A. Proteins are categorized as ‘redundant’ if they have only shared peptides that can be assigned to other entries and thus these proteins are not needed to explain the observed peptide evidence (tiers 7-9). Finally, all other proteins that completely lack any peptides observed at our minimum PSM significance threshold are categorized as ‘not observed’ (tier 10).

### Handling of gene models and splice forms

The 27655 protein coding genes in Araport11 are represented by 48359 gene models (transcript isoforms), which are identified by the digit after the AT identifier (*e.g.* AT1G10000.1). We refer to the translations of these gene models as protein isoforms. Most protein isoforms are very similar (differing only a few amino acid residues often at the N- or C-terminus) or even identical at the protein level. It is often hard to distinguish between different protein isoforms due to the incomplete sequence coverage inherent to most MS proteomics workflows. For the assignment of canonical proteins (at least two uniquely mapping peptides identified) (Table 2A,B), we selected by default only one of the protein isoforms as the canonical protein; this was the ‘.1’ isoform unless one of the other isoforms had a higher number of matched peptides. However, if other protein isoforms did have detected peptides that are unique from the canonical protein isoform (*e.g.* perhaps due to a different exon), then they can be given tier 1 or less confident tier status depending on the nature of the additional uniquely mapping peptides (length and numbers) (Table 2A,B). If the other protein isoforms do not have any uniquely mapping peptides amongst all protein isoforms (for that gene), then they are classified as redundant (tiers 7-9 in the more complex system).

### Protein physicochemical properties and functions

To characterize the canonical and unobserved proteomes, physicochemical properties were calculated or predicted using various web-based tools. These include: protein length, mass, GRAVY index, isoelectric point (pI), number of transmembrane domains (http://www.cbs.dtu.dk/services/TMHMM) and sorting sequences for the ER, plastids and mitochondria (http://www.cbs.dtu.dk/services/TargetP-1.0/).

### Integration of PeptideAtlas results in other web-bases resources

PeptideAtlas is accessible through its web interface at http://peptideatlas.org. Furthermore, direct links are provided between PeptideAtlas and PPDB (http://ppdb.tc.cornell.edu/), UniProtKB (https://www.uniprot.org/) and TAIR (https://www.arabidopsis.org/) at the level of protein entries. Links to matched peptide entries in PeptideAtlas are available in the Arabidopsis annotated genome through a specific track in JBrowse at https://jbrowse.arabidopsis.org.

## RESULTS AND DISCUSSION

### Overview of the generation and output of the Arabidopsis PeptideAtlas

Figure 1 provides an overview of the generation of the first build of the Arabidopsis PeptideAtlas. The project started by collecting all available MS datasets for Arabidopsis from ProteomeXchange; we refer to these datasets as PXDs. A subset of PXDs was selected (see next section) and detailed information about the samples and MS acquisition within each PXD was collected and annotated using a newly built in-house meta-data annotation system. Selected PXDs were processed through the TPP to match MS data to peptides and proteins (including selected PTMs) in Araport11, TAIR10, a collection of small peptides, as well as other predicted proteins (Table 1). The genome annotation of Araport11 was used as default (see Methods). For each analyzed PXD, we calculated the MSMS spectral match rate to peptides as a measure for MSMS data quality as well as data processing. In case of very low match rate (<10%) we reevaluated the search parameters and, if needed, reran the search with adjusted parameters. Following rigorous evaluation using sophisticated dedicated algorithms to control false discovery rates and PTM site verification (Shteynberg et al., 2019), as well redundancy removal (avoiding identical predicted proteins listed under different protein identifiers), identified proteins were classified into a ten-tier system, ranging from very high confidence identifications to low confidence identifications (Table 2A). We also provide a simpler four-category system in which tiers 2-7 are folded into a single category (Table 2B). The identified proteome was then evaluated for physicochemical properties, predicted subcellular localization and function. This first Arabidopsis PeptideAtlas build is freely available and protein entries are directly linked to TAIR, PPDB and UniProtKB. Peptides are mapped to the Arabidopsis genome on specific tracks through the genome browser JBrowse. After in-depth evaluation of the identified proteome coverage from this first build and feedback from the international research community, we will select additional PXDs for subsequent PeptideAtlas builds, as discussed further below in the section ‘*The next Arabidopsis PeptideAtlas build*’. In the remainder of this paper we will provide more detail about and insights into this PeptideAtlas build and the observed Arabidopsis proteome.

**Figure 1.**
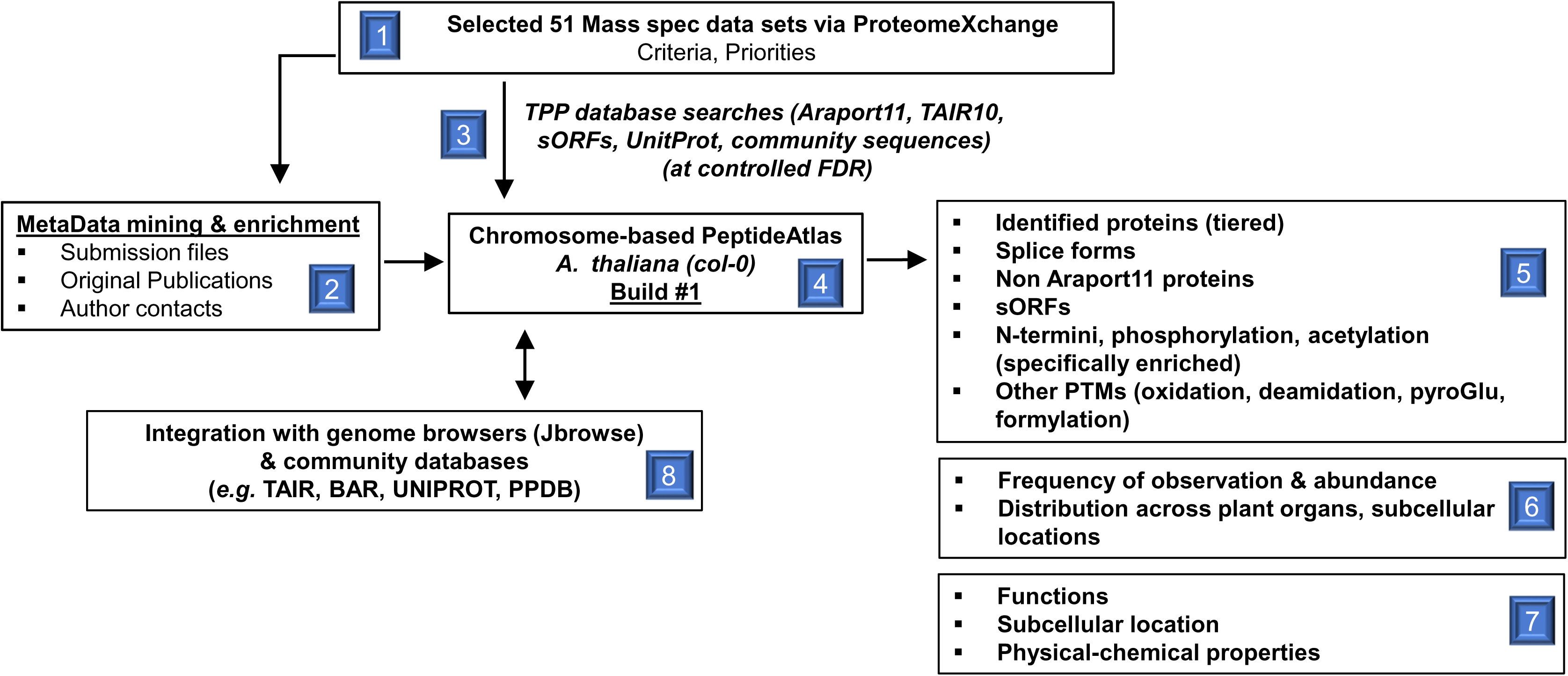
Graphical overview of the Arabidopsis PeptideAtlas project and generation of the first build presented here. Specific steps and components are numbered.

### Features of publicly available Arabidopsis PXDs

At the start of building of this first Arabidopsis PeptideAtlas in the fall of 2019, we first reviewed all PXDs available through the ProteomeXchange interface for Arabidopsis, and we continued to so do as the project progressed into 2020. We verified if indeed the plant material was *Arabidopsis* (and also checked the ecotype), scored each submission for the type of MS instrument(s) with which the data were collected, and collected information about nature of the samples (*e.g*. organ, subcellular fraction, enrichment for specific PTMs). Figure 2 summarizes some of this information for all 356 Arabidopsis PXDs until 15 July 2020. The first Arabidopsis PXD available in ProteomeXchange was from 2012 (we note that earlier submissions to PRIDE (Perez-Riverol et al., 2018) were not transferred to ProteomeXchange) and the number of datasets exponentially increased in subsequent years, resulting in 357 available PXDs from some 200 different laboratories by July 2020 (Figure 2A). A wide range of MS instruments were used to acquire these data (Figure 2B). There were just four submissions that used MALDI-TOF-TOF instruments and the majority (82%) used different generations of Orbitrap-based instruments from Thermo Fisher Scientific (Eliuk and Makarov, 2015; Makarov, 2019). The sensitivity and throughput of MS has dramatically increased over subsequent generations of MS instruments and this should also be reflected in increasing proteome coverage with newer PXDs. Based on keywords and information associated with each PXD, figure 2C gives an impression of the types of subcellular fractions analyzed across these 356 PXDs. For simplicity we grouped various keywords into 10 sample types, which showed a strong interest in proteomes from chloroplasts (often specific sub-organellar fractions such as thylakoids, stroma or enveloped). It should be noted that many of the PXDs did not involve a specific subcellular fraction, but rather analyzed proteome extracts (either just soluble or total detergent extracted proteomes) from whole seedlings, plant parts (*e.g*. roots, flowers, rosettes) without further subfractionation. Whereas all PXDs allowed for one or more common PTMs that are either (often) induced after protein extraction (*e.g.* oxidation of Met or Trp, cyclization of Gln and Glu, deamidation of Asn or Gln, and carbamidomethylation of Cys), a subset of PXDs specifically focused on selected PTMs that often require affinity enrichment or labelling (Figure 2D). A significant portion of PXDs focused on protein phosphorylation, N-terminal or lysine acetylation, ubiquitination and various cysteine modifications. Finally, these proteomics analyses were motivated by a wide range of biological questions, including abiotic stress (*e.g.* cold, heat, light, oxidation, metals, touch), biotic stress/plant immunity (*e.g. Pseudomonas syringae*, flagellin), developmental questions (*e.g.* seed development and germination) and circadian rhythm.

**Figure 2.**
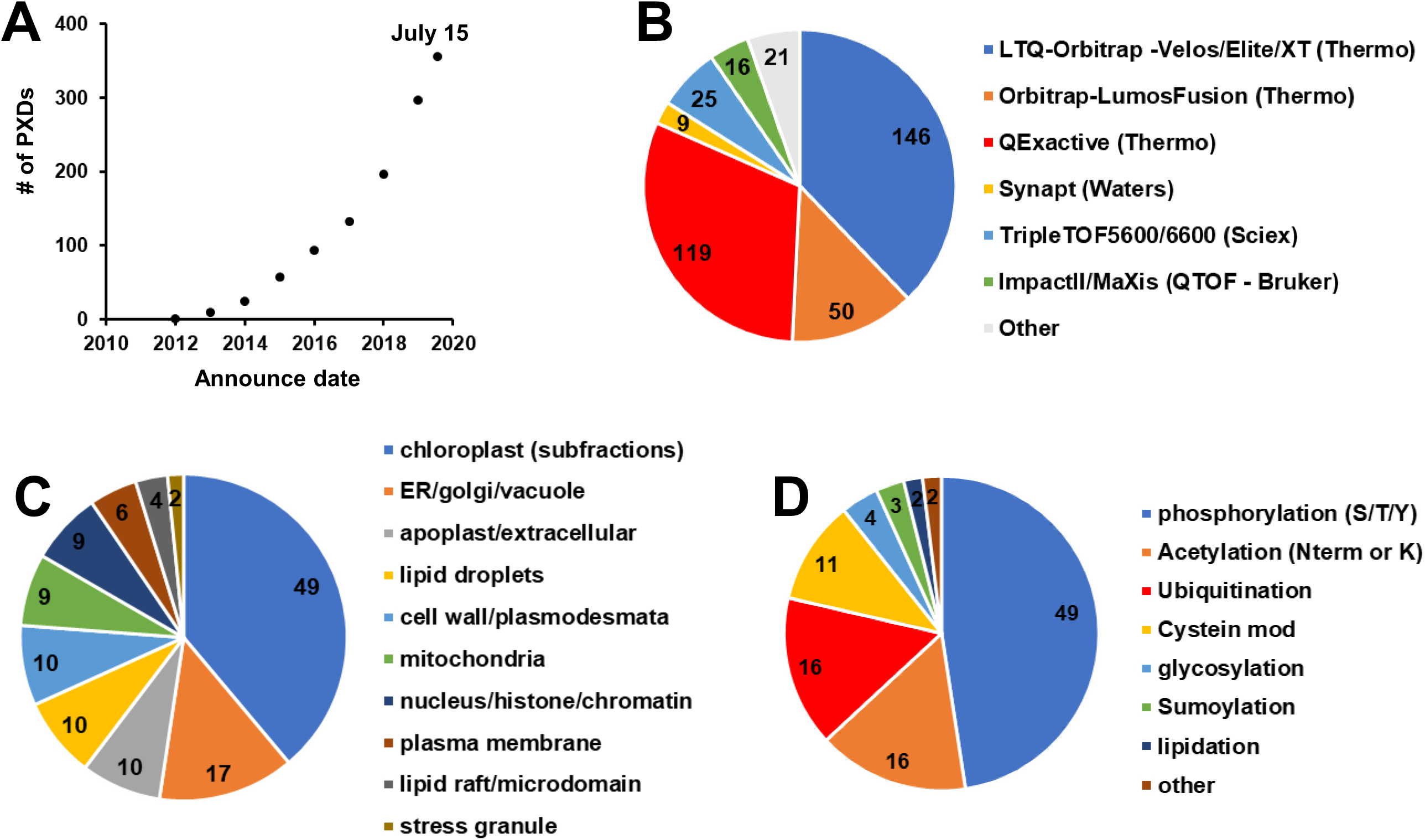
Features of PXDs for Arabidopsis available via ProteomeXchange through July 15, 2020. Information for these PXDs was obtained from the submitted metadata and/or accompanying publications. **(A)** Accumulative PXDs with verified Arabidopsis content by year (2010-7/2020) **(B)** Type of MS Instrument (LTQ-Orbitrap - Velos/Elite/XT (Thermo), Orbitrap-LumosFusion (Thermo), QExactive (Thermo), Synapt (Waters), TripleTOF5600/6600 (Sciex), ImpactII/MaXis (Bruker), other. **(C)** Arabidopsis subcellular fractions (plastid, mitochondria, peroxisomes, vacuole, nucleus, apoplast/extracellular, cytosol, ER/Golgi/PM) **(D)** Post-translational modifications that were specifically enriched prior to MS analysis (phosphorylation, acetylation (N-term or Lys), ubiquitination, cysteine oxidation, glycosylation, sumoylation, lipidation, other).

### Selection of PXDs for the first build

Because it was not feasible to process all available PXDs for this first PeptideAtlas build (due to time and computing constraints), we focused mostly on those PXDs that were generated by the high mass accuracy Orbitrap-type instrument types since they were by far the most frequently used (Figure 2B) and to simplify the data analysis and better control false discovery rates. Table 3 provides key information for the final 52 PXDs used in this first build; additional details are provided in Supplemental Dataset 1. The majority of selected PXDs were from 2019 (∼40%), with additional PXDs from 2015-2018. We also added the recent 2020 PXD (PXD013868) associated with (Mergner et al., 2020) because it included a very large amount of MS data, including phosphorylated proteins, sampled across 30 different Arabidopsis tissues. Finally, most PXDs were from ecotype Colombia 0 (Col-0) since this is the reference Arabidopsis ecotype that was originally sequenced and on which the *Arabidopsis* Araport11 and previous TAIR genome annotations are based. However, one PXD used *Wassilewskij* and several PXDs used ecotype *Landsberg erecta* mostly for cell cultures (ordered from the Arabidopsis Biological Resource Center (ABRC); PSB-D (CCL84840) and PSB-L (CCL84841)).

**Table 3.** Summarizing information of the 52 selected PXD datasets for this first PeptideAtlas build. This includes PXD #, publication, number of matched MSMS spectra and % match rate, the number of identified proteins (canonical and groups of proteins), the number of matched distinct MSMS peptides, the MS instrument, information about the sample (plant part, subcellular fraction, enrichment for PTMs. An extended table with additional information is provided as Supplemental Dataset 1.

| Dataset Identifier | Publication | matched # of MS/MS spectra | matched MS/MS spectra (%) | # Distinct peptides | Instrument | Plant Parts | Subcellular Fraction | N-terminome & specific PTM analysis |
| --- | --- | --- | --- | --- | --- | --- | --- | --- |
| PXD000136 | Hesse et al. (2016) | 22,419 | 0.16 | 4,569 | LTQ FT | rosette leaves | chloroplast |  |
| PXD000546 | Tomizoli et al. (2014) | 120,945 | 0.43 | 8,858 | LTQ Orbitrap Velos | rosette leaves | chloroplast |  |
| PXD000660 | Köhler et al. (2015) | 11,795 | 0.12 | 3,280 | LTQ Orbitrap Velos | rosette leaves | chloroplast | N-terminome (TAILS) |
| PXD000869 | Zhang et al. (2018) | 62,763 | 0.40 | 4,616 | LTQ Orbitrap Velos | rosette leaves | chloroplast |  |
| PXD000908 | Baerenfaller et al. (2015) | 466,846 | 0.22 | 19,973 | LTQ Orbitrap XL | rosette leaves |  |  |
| PXD001207 | Köhler et al. (2015) | 26,393 | 0.35 | 7,554 | LTQ Orbitrap Velos | rosette leaves | chloroplast |  |
| PXD001473 | Li-Ling Lin et al. (2015) | 13,313 | 0.08 | 655 | LTQ Orbitrap XL | cell culture (ecotype Landsberg erecta) |  | phosphorylation |
| PXD001719 | Zhang et al. (2015) | 39,966 | 0.20 | 13,154 | LTQ Orbitrap Velos | root |  | N-terminome (TAILS) |
| PXD001855 | Venne et al. (2015) | 36,078 | 0.12 | 12,721 | Q Exactive | seedling |  | N-terminome (ChaFRADIC) |
| PXD002069 | Linster et al. (2015) | 213,875 | 0.08 | 7,230 | LTQ Orbitrap Velos | rosette leaves |  | acetylation of N-term and lysine |
| PXD002160 | Correa-Galvis et al. (2016) | 77,605 | 0.14 | 3,638 | LTQ Orbitrap Elite | rosette leaves | chloroplast |  |
| PXD002186 | Nishimura et al. (2015) | 257,800 | 0.45 | 9,575 | LTQ Orbitrap | rosette leaves | chloroplast |  |
| PXD003162 | Lundquist et al. (2017) | 247,024 | 0.22 | 13,092 | LTQ Orbitrap Elite | rosette leaves | chloroplast |  |
| PXD003516 | Wang et al. (2016) | 25,046 | 0.17 | 12,277 | Q Exactive | rosette leaves | chloroplast |  |
| PXD003684 | Bhuiyan et al. (2016) | 130,083 | 0.32 | 8,337 | LTQ Orbitrap | rosette leaves | chloroplast |  |
| PXD004025 | Al Shw eiki et al. (2017) | 559,931 | 0.39 | 22,301 | LTQ Orbitrap Velos | rosette leaves |  |  |
| PXD004276 | Choudhary et al. (2016) | 59,548 | 0.19 | 12,591 | LTQ Orbitrap | seedling |  | phosphorylation |
| PXD004599 | Mattei et al. (2016) | 11,582 | 0.13 | 2,371 | LTQ Orbitrap | seedling |  | phosphorylation |
| PXD004742 | Subramanian et al. (2016) | 161,009 | 0.45 | 8,788 | LTQ Orbitrap Velos | rosette leaves |  |  |
| PXD004896 | Willems et al. (2017) | 87,053 | 0.13 | 31,022 | LTQ Orbitrap | cell culture (ecotype Landsberg erecta) |  | N-terminome (COFRADIC) |
| PXD005600 | Schonberg et al. (2017) | 45,886 | 0.13 | 2,462 | LTQ Orbitrap Velos | rosette leaves | chloroplast | phosphorylation |
| PXD005740 | Hander et al. (2019) | 1,388 | 0.02 (a) | 860 | Q exactive | seedlings (split in root & rosette) |  |  |
| PXD006113 | Brocard et al. (2017) | 153,376 | 0.23 | 13,578 | LTQ Orbitrap | rosette leaves | lipid droplet |  |
| PXD006328 | Strehmel et al. (2017) | 32,278 | 0.10 | 6,268 | Q Exactive | root | exudate |  |
| PXD006347 | Nee et al. (2017) | 9,155 | 0.03 (b) | 1,838 | Q Exactive | seed |  |  |
| PXD006651 | Hartl et al. (2017) | 159,852 | 0.61 | 26,626 | Q Exactive | leaf/rosette | chloroplast | lysine acetylation |
| PXD006652 | Hartl et al. (2017) | 114,752 | 0.25 | 15,734 | Q Exactive | leaf/rosette | chloroplast | lysine acetylation |
| PXD006800 | Brault et al. (2019) | 238,014 | 0.47 | 28,668 | Q Exactive | cell culture (ecotype Landsberg erecta) | total cell extract, plasmodesmata, plasma membrane, microsome & cell wall |  |
| PXD006806 | Brault et al. (2019) | 634,158 | 0.73 | 40,233 | Q Exactive | cell culture (ecotype Landsberg erecta) | plasmodesmata, plasma membrane, microsome & cell wall |  |
| PXD006848 | Seaton et al. (2018) | 1,609,008 | 0.54 | 51,874 | LTQ Orbitrap Velos | rosette leaves |  |  |
| PXD007630 | Koskela et al. (2018) | 166,166 | 0.35 | 15,184 | Q Exactive | rosette leaves | chloroplast | N-terminal/lysine acetylation |
| PXD008355 | Van Leene et al. (2019) | 374,427 | 0.28 | 21,350 | Q Exactive | cell culture (ecotype Landsberg erecta) |  | phosphorylation |
| PXD008663 | Castrec et al. (2018) | 172,302 | 0.05 (c) | 6,906 | LTQ Orbitrap Velos | rosette leaves |  | N-term & lysine acetylation |
| PXD009016 | Zhang et al. (2019) | 94,355 | 0.15 | 13,443 | Q Exactive | rosette leaves |  | phosphorylation |
| PXD010324 | Waltz et al. (2019) | 402,558 | 0.31 | 17,491 | Q Exactive | flow ers; cell culture (dark) | mitochondria |  |
| PXD010545 | Bouchnak et al. (2019) | 66,782 | 0.19 | 16,619 | Q Exactive | rosette leaves (ecotype Wassilew skja) | chloroplast |  |
| PXD010730 | Wu et al. (2019) | 629,552 | 0.44 | 27,754 | Q Exactive | rosette leaves |  |  |
| PXD011088 | Rugen et al. (2019) | 589,487 | 0.23 | 22,692 | Q Exactive | rosette leaves; cell culture (Col-0) | mitochondria |  |
| PXD011483 | McLoughlin et al. (2019) | 2,897,014 | 0.38 | 44,923 | Q Exactive | rosette leaves; seedlings | protein aggregates |  |
| PXD011716 | Kosmacz et al. (2019) | 121,240 | 0.17 | 24,030 | Q Exactive | seedlings | stress granule |  |
| PXD011759 | Wu et al. (2019) | 840,034 | 0.45 | 40,953 | Q Exactive | seedlings |  |  |
| PXD012708 | Zhang et al. (2019) | 6,316,858 | 0.47 | 239,706 | Orbitrap Fusion Lumos | 10 plant parts (rosette leaves, cauline leaf, stems, flow er, pollen, siliques, seeds, cotyledons, root, root cell culture) |  |  |
| PXD012710 | Zhang et al. (2019) | 2,077,659 | 0.15 | 123,441 | TripleTOF 5600 | 11 plant parts (rosette leaves, cauline leaf, stems, flow er, pollen, siliques, seeds, cotyledons, root, root cell culture) |  |  |
| PXD013005 | Wu et al. (2019) | 731,387 | 0.35 | 41,004 | Q Exactive | seedlings |  |  |
| PXD013325 | Jiang et al. (2019) | 15,139 | 0.23 | 4,558 | LTQ Orbitrap Elite | rosette leaves |  |  |
| PXD013494 | Montandon et al. (2019) | 29,810 | 0.19 | 3,782 | LTQ Orbitrap | rosette leaves | chloroplast |  |
| PXD013637 | Hu et al. (2019) | 77,866 | 0.14 | 15,379 | Q Exactive | rosette leaves |  |  |
| PXD013646 | Furtauer et al. (2019) | 2,374,645 | 0.21 | 35,719 | Q Exactive; LTQ Orbitrap Elite | rosette leaves (ecotype Landsberg erecta) |  | phosphorylation |
| PXD013868 | Mergner et al. (2020) | 15,180,331 | 0.27 | 388,665 | Q Exactive HF | 30 tissue types |  | phosphorylation |
| PXD017189 | Bhuiyan et al. (2020) | 59,880 | 0.31 | 5,203 | LTQ Orbitrap | rosette leaves |  |  |
| PXD017380 | not published | 267,019 | 0.12 | 24,271 | Q Exactive | rosette leaves | chloroplast |  |
| PXD017400 | not published | 367,359 | 0.17 | 19,279 | Q Exactive | rosette leaves | chloroplast |  |
| **Totals** |  | **39,480,811** |  |  |  |  |  |  |

(a) Affinity pulldown GFP tagged protein
(b) Native peptides (peptidome); no digests
(c) N-acetoxy- $[\text{H}_3]$ succinimide ( $\text{C}_6\text{D}_5\text{H}_4\text{NO}_4$ ) treatment labeling resulting N-terminal and Lys acetylation as well as O-acetylation of Ser, Thr, and Tyr side-chains but then reversed through hydrolysis

We aimed to have representation across as many plant parts as possible to maximize proteome coverage, including those proteins that are specifically expressed in specific parts of the plants (see Table 3 and Figure 3). Figure 3A shows the number of MS runs for the different types of plant samples. The vast majority of MS runs (61%) were done on the main green tissues, including whole rosettes, specific leaf stages, cauline leaves, stems and petioles. 14% of the MS runs were done on whole siliques, seeds in different developmental stages or embryos isolated from seeds. Root samples (tip, whole roots or even root exudates) were analyzed in 7.4% of the MS runs, whereas whole flowers or specific flower parts were used in 5% of the MS runs. Cell cultures were used in 6.6% of the MS runs. Finally, a smaller number of MS runs (0.5-1.4%) were from hypocotyls, callus, pollen, cotyledons or young seedlings (including roots, cotyledons and a few leaves). For most of these MS runs, there was no further subcellular fractionation, and the proteome was either extracted in the presence of the strong ionic detergent SDS or in the absence of detergent, resulting in the extracted total cellular proteome including membrane proteins or just the soluble proteome, respectively. However, for nearly 20% of the MS runs, subcellular fractions were isolated from the plant parts, in particular isolated chloroplasts or sub-chloroplast compartments (thylakoids, stroma, envelopes, nucleoids, or plastoglobules), but also mitochondrial fractions (mostly ribosomes) (Figure 3B). Other subcellular fractions included cytosolic lipid droplets, cytosolic stress granules, root exudate and enriched plasmodesmata fractions (Figure 3B). The relative high number of chloroplast samples is because the proteomes of chloroplasts have been the subject of many of the PXDs over the last 10 years (Figure 2C), and also because of our own expertise and interests in chloroplasts.

**Figure 3.**
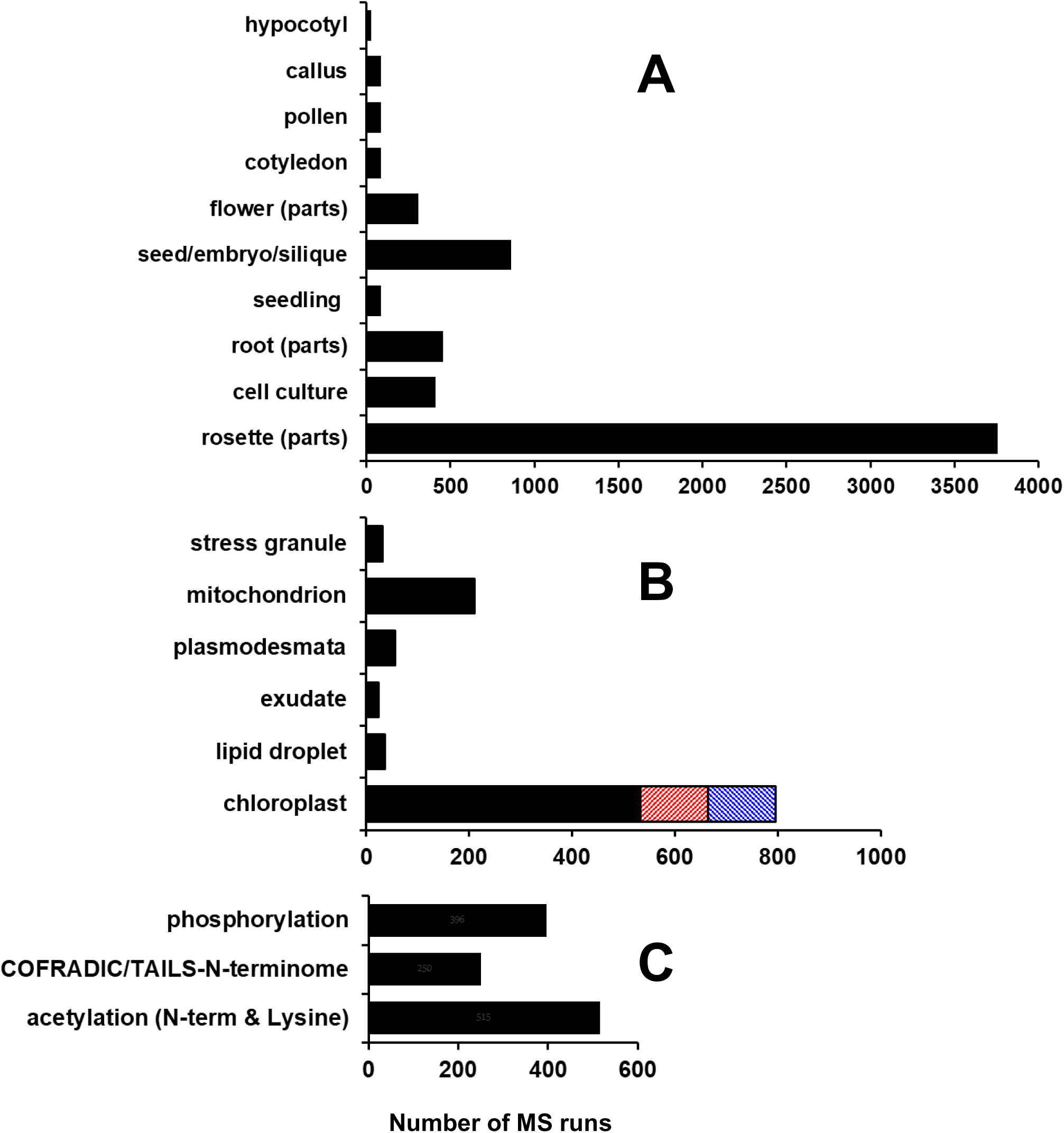
Key features of samples used for the raw files (MS runs) for the 52 selected PXDs for this first Arabidopsis PeptideAtlas build. The count is based on the number of MS runs (raw files) for each part. **(A)** Arabidopsis plant parts – hypocotyl, callus, pollen, cotyledon, flower parts (sepal/petal/carpel/stamen/pedicle), seed/septum/embryo), seedling (all parts of a young plant - root-hypocotyl/cotyledons/few young leaves, mostly collected from plates or liquid culture, root (tip/exudate/zone), cell culture, rosette parts (rosette/leaf/petiole/cauline leaf/senescing leaf/stem/internode). **(B)** Arabidopsis subcellular fractions specifically analyzed are stress granule, mitochondrion, plasmodesmata, root exudate, cytosolic lipid droplet, chloroplast (black) and the specific fractions thylakoid (orange), plastoglobuli (blue). **(C)** MS runs of samples that were specifically prepared to analyze PTMs (phosphorylation, acetylation of the N-terminus and/or lysine) or determine the physiological N-terminus using N-terminome enrichment techniques (TAILS, COFRADIC or ChaFRADIC).

To support recognition and annotation of the N-termini of mature proteins (including after maturation processes such as cleavage of signal peptides), we selected several PXDs in which specific N-terminal labeling and enrichment techniques (TAILS (Marino et al., 2015); COFRADIC (Staes et al., 2011)) were used to identify the N-termini of accumulated proteins, protein-derived signaling peptides or protein degradation products (Figure 3C). Finally, the set of PXDs also included the most studied PTMs *i.e.* phosphorylation and N-terminal or lysine acetylation (Figure 3C).

### The identified proteome in the first PeptideAtlas build and MS support

MS data can realistically (avoiding *de novo* annotation of MSMS spectra) only lead to identification of peptides and proteins by searching these MS data against an assembly of predicted, putative proteins. Proteins or peptides not represented in this protein search space cannot be identified. Therefore, we assembled a comprehensive set of sequences from a variety of key sources (Table 1). These are Araport11 (the most recent (2017) annotation of the Arabidopsis genome), TAIR10 as the precursor of Araport11, RefSeq and UniProtKB, a large collection of putative and speculative peptides encoded by sORFs (assembled in ARA-PEP - (Hazarika et al., 2017)), as well as a collection of high expression putative orphan ORFs from E. Wurtele (Iowa State). This totaled 204,154 sequence identifiers with significant redundancy and smaller numbers of unique proteins for each source (see Table 1); overall these represented 73,816 unique amino acid sequences. Following downloading of PXD raw MS files, file conversions and sample annotations, the MS data were searched against this total protein space (see METHODS). We searched in several iterations to optimize the search parameters (mostly variable and fixed PTMs) and search time. In particular PXDs involving stable isotope dimethylation for N-terminomics and lysine acetylation (see Table 3) required particular attention, because these can lead to different mass shifts in dependence of the isotopes employed (+28 (2x C^12^H_3_) for light; +32 (2xC^12^HD_2_) or +34 (2x C^12^D_3_ or C^13^HD_2_) for heavy). Also the use of tandem mass tag (TMT) or isobaric tags for relative and absolute quantitation (iTRAQ) labeling used for multiplexing and comparative proteomics required careful attention and verification of metadata.

The finalized searches and post-search processing for control of false discovery rates resulted in the matching of nearly 40 million out of about 143 million submitted MSMS spectra, leading to the identification of 535340 distinct peptides matching to 17858 canonical proteins, as well as 1942 uncertain and 1600 redundant proteins for which identification is ambiguous due to shared peptides or lower evidence levels (http://www.peptideatlas.org/builds/arabidopsis/). For the remaining 6255 proteins there were no observed matching peptides (Table 3; Supplemental Dataset 1). The overall match rate of MSMS spectra to peptides was 28%, but this match rate varied dramatically across PXDs from 2% to 74% (Table 3), with an average and median match rate of 26% and 22%, respectively. For those PXDs where we obtained a low match rate, we re-evaluated the search parameters to ensure that we did not overlook specific sample treatments that could affect the optimal search parameters (*e.g*. labeling techniques). The low match rate (<10%) was in most cases observed for N-terminomics and acetylation (N-terminal and lysine) studies involving dimethyl-labeling possibly combined with TAILS or COFRADIC/ChaFRADIC and in other cases involving affinity purification with a specific bait or analysis of the secreted peptidome from roots (Table 3). Other explanations for variations in match rate are often related to the acquisition parameters in particular low thresholds for MSMS acquisition and/or lack of repeat MSMS scans, resulting in low quality MSMS spectra. We did not detect an obvious relationship between MSMS match rate and instrument type across the PXDs.

Figure 4 shows the number of distinct (non-redundant) peptides (irrespective of PTMs) (Figure 4A) and distinct identified canonical proteins (Figure 4B) as function of the cumulative number of matched MSMS spectra ordered by PXD identifier (from low to high or old to new) for this first Arabidopsis PeptideAtlas. To better understand the underlying data for this PeptideAtlas build, we calculated the frequency distributions of peptide charge state, missed cleavages and peptide length for the ∼40 million matched MSMS spectra (Figure 5). The vast majority of matched spectra had a charge state of +2 (60%), +3 (34%) or 4+ (5.6%) and minor amounts of 1+ (0.09%), 5+ (0.71%) or 6+ (0.08%) (Figure 5A). The majority of matched tryptic PSMs (77%) did not have a missed cleavage, whereas 20%, 3% and 0.1% had 1, 2 or 3 missed cleavages, respectively (Figure 5B). Allowing for missed cleavages can potentially increase the false peptide discovery rate because it increases the peptide search space, but it is not uncommon that missed cleavages occur and it does allow for increased sequence coverage and detection of N- and C-termini and splice junctions. We observed a wide range of matched peptide lengths, with seven amino acids being the shortest sequence allowed (Figure 5C). 99% of all matched peptides were between 7 and 35 aa length, with the most frequent peptide length of 12 aa.

**Figure 4.**
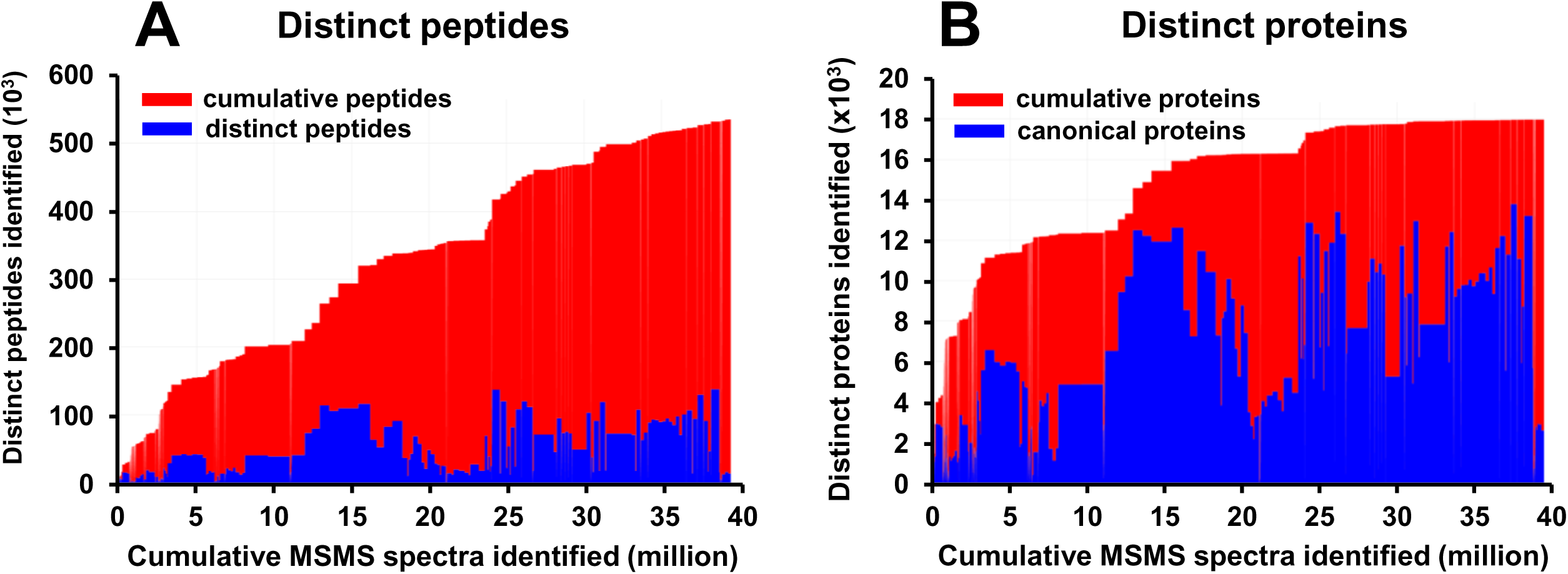
Number of distinct (non-redundant) peptides (left panel) and identified canonical proteins (right panel) as a function of the cumulative number of PSMs (peptide-spectrum matches) for the first Arabidopsis PeptideAtlas. The cumulative count is ordered by PXD identifier (from low to high or old to new). The build is based on 266 experiments across the 52 selected PXDs, where each PXD may be decomposed into several experiments/samples when such information can be determined). The PSM FDR is 0.0001. **(A)** Number of distinct (non-redundant) peptides as function of the cumulative number of MSMS spectra matched. 535,000 distinct peptides are identified at a peptide-level FDR of 0.001. Areas in blue indicate the total number of distinct peptides in each experiment, whereas areas in red indicate the cumulative number of identified peptides from the current and previous experiments. **(B)** Number of distinct (non-redundant) canonical proteins as function of the cumulative number of MSMS spectra matched at a canonical protein-level FDR of 0.0005. Areas in blue indicate the total number of distinct canonical proteins in each experiment, whereas area in red indicate the cumulative number of identified canonical proteins from the current and previous experiments.

**Figure 5.**
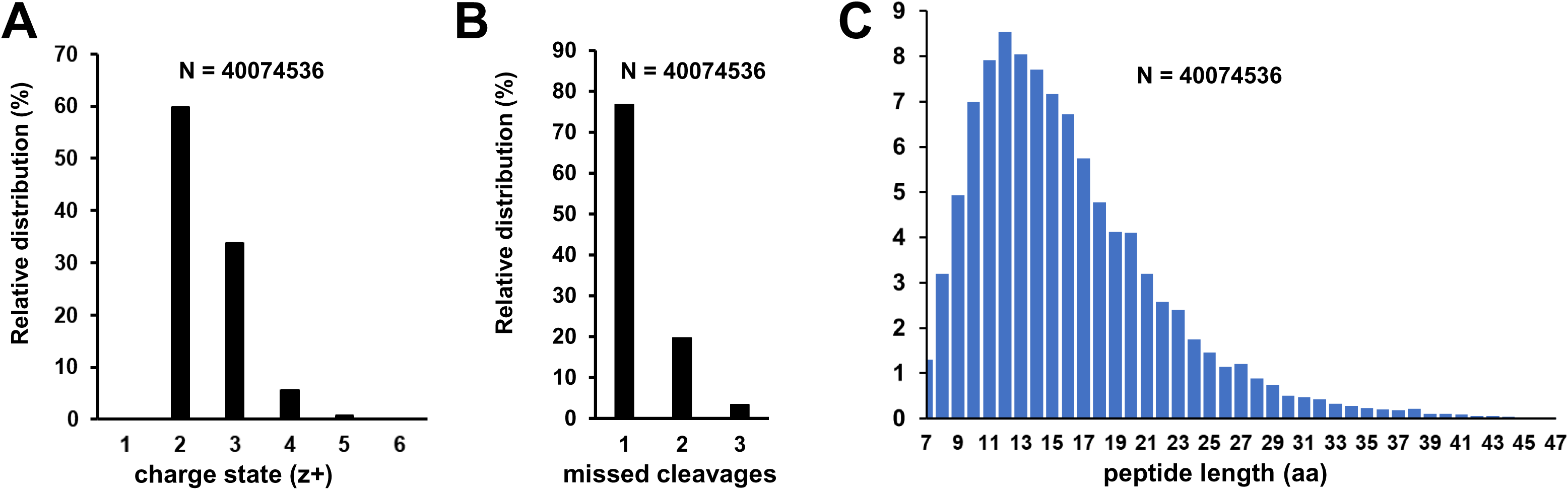
Key statistics of matched MSMS data for this PeptideAtlas build. **(A)** Frequency distribution of peptide charge state (z) **(B)** Frequency distribution missed cleavages for tryptic peptides. Note that when R or K is followed by P, trypsin does not cleave and hence these are not counted towards missed cleavages. **(C)** Frequency distribution of peptide length (aa)

#### Mapping the Araport11 proteome and splice forms

Because the Araport11 annotation is the most common reference used by the Arabidopsis community compared to TAIR10, RefSeq and UniProtKB, the default protein identifier for sets of identical protein sequences (across all sources) was always from Araport11. Araport11 has 27655 protein coding genes with 48359 gene model or transcript isoforms (Cheng et al., 2017), representing 40784 unique protein amino acid sequences; it should be noted that the difference between transcript isoforms for a gene are often very minor at the amino acid level. For comparison, TAIR10 has 27416 genes and 35386 transcript isoforms, representing 32785 unique proteins; 1651 protein sequences are found TAIR10 but not in Araport11 (at 100% sequence identity) (Table 1). The vast majority of peptide sequences (>99%) in this first build matched to proteins in Araport11 (Table 4) with the remainder matching to sequences in one or more of the other sources (Table 5). We assigned multiple confidence levels of protein identification using a sophisticated tiered system (with ten tiers) similar as was developed for the human PeptideAtlas (Deutsch et al., 2016a) (Table 2A). These ten tiers allow to precisely distinguish different evidence levels of protein identification, including the use of peptides that are matched to multiple proteins (see METHODS). Figure 6A shows a schematic explanation for the tier system and figures 6B-D provide specific examples from this PeptideAtlas build. These ten tier assignments were then also condensed in a simplified classification of proteins identification with just four categories (Table 2B) to provide an easier overview of the identified proteome. The overall number of identified proteins for both classifications systems is displayed in the PeptideAtlas browser. (http://www.peptideatlas.org/builds/arabidopsis/).

**Figure 6.**
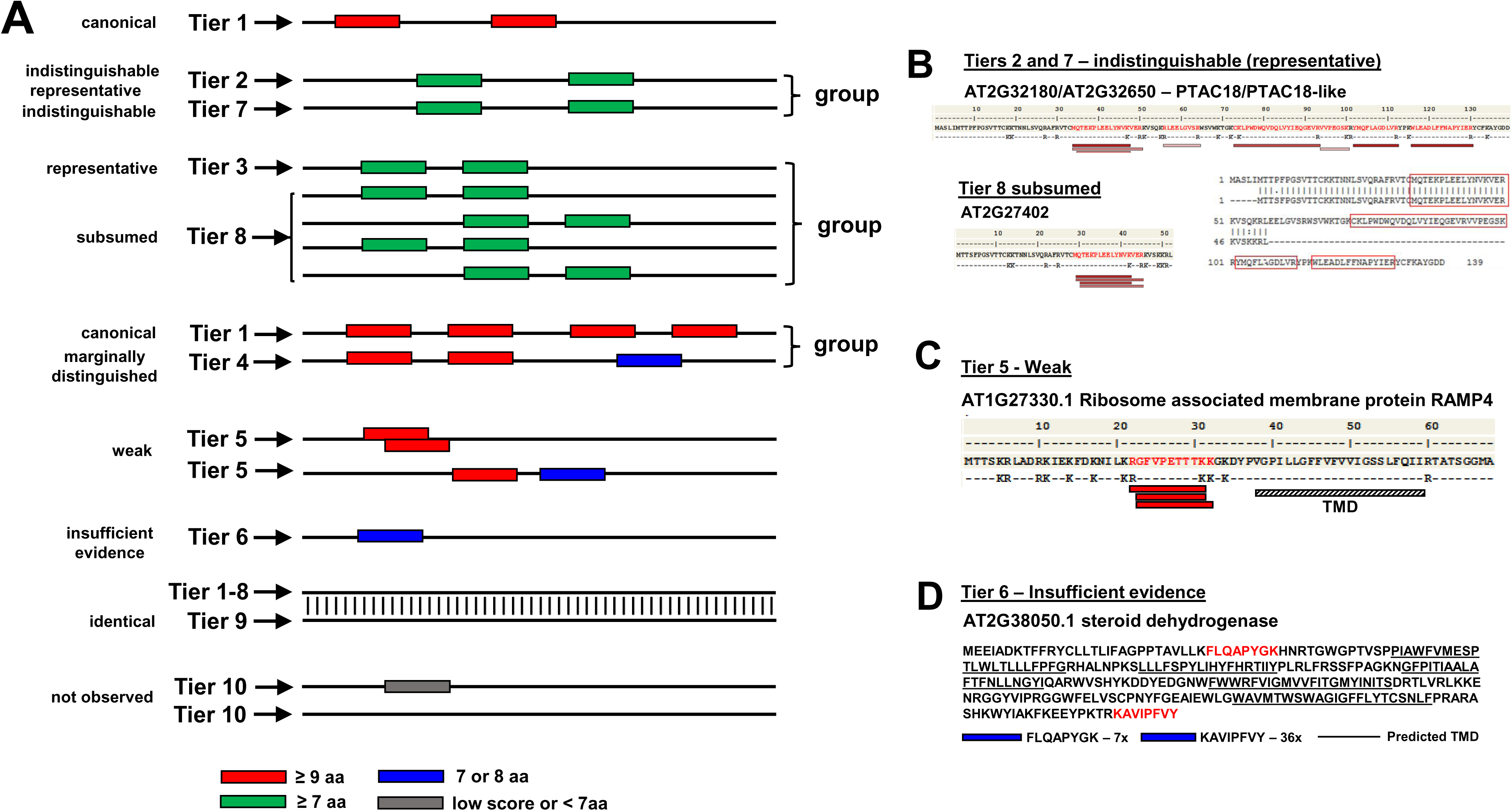
Explanation and examples of the tiered identification system. **(A)** Schematic depiction of the tiered protein identification system. Protein sequences are represented by a simple line and identified peptides (PSMs) are shown as filled rectangles of different color. Peptides contributing to identification at the highest confidence level (tier 1 – canonical) are shown in red (must be at least 9 amino acids). Peptide of 7 or more amino acids are shown in green. Peptides of 7 or 8 amino acids are shown in blue. Peptides less than seven amino acid are never considered for protein identification and are shown in grey. Also any PSMs below a minimum build threshold of 0.001 PSM-level FDR are shown in grey. This panel shows eight scenarios where either a single protein is identified or where a group of proteins in identified. **(B)** This panel shows a case where three proteins were identified in a group. Two identical proteins AT2G32180 (PTAC8) and AT2G32650 (PTAC18-like) were identified with nine distinct peptides. Because these proteins are identical in sequence one cannot distinguish them, one was designated the indistinguishable representative (tier 2) and the other one as indistinguishable (tier 7). A third protein AT2G27402 with partial sequency identity was identified by a subset of these distinct peptides and was therefore assigned to tier 8 (subsumed) because this protein is not needed to explain these PSMs. An aa alignment between PTAC8/PTAC8-Like and AT2G27402 shows the residues that were part of the identified peptides (boxed in red). **(C).** This panel shows an example of AT1G27330.1 of a tier 5 identification (weak). This is a small Ribosome Associated Membrane Protein (RAMP4) (68 aa) with one predicted transmembrane domain in the C-terminal portion, a positive GRAVY index (0.034) and three nested or overlapping peptides, each identified multiple times across several independent PXD datasets and publications. Moreover, the N-terminal region contains eight closely spaced lysine and arginine residues which would generate very short (3-5 aa) peptides that are too small to be considered as supported evidence by MSMS. **(D)** Figure 6D shows a tier 6 example of a steroid dehydrogenase (ATDET2/ DWARF6; AT2G38050.1) involved in the brassinolide biosynthetic pathway. It has five or six predicted transmembrane domains and a positive GRAVY index of 0.132. It was identified in two publications across some 40 different sample types with a nine aa N-terminal peptide (just downstream of a hydrophobic region) and an eight aa C-terminal peptide.

**Table 4.** Proteins identified in Araport11 for each of the four confidence categories by nuclear chromosome (1 to 5), mitochondrial (M) and plastid chromosome (C).

| Chromosome | Entries | Canonical |  | Uncertain |  | Redundant |  | Not Observed |  |
| --- | --- | --- | --- | --- | --- | --- | --- | --- | --- |
| **M** | 122 | 15 | 12.3% | 10 | 8.2% | 17 | 13.9% | 80 | 65.6% |
| **C** | 88 | 59 | 67.0% | 15 | 17.0% | 7 | 8.0% | 7 | 8.0% |
| **1** | 7156 | 4622 | 64.6% | 545 | 7.6% | 397 | 5.5% | 1592 | 22.2% |
| **2** | 4317 | 2695 | 62.4% | 291 | 6.8% | 247 | 5.7% | 1084 | 25.1% |
| **3** | 5460 | 3561 | 65.2% | 365 | 6.7% | 308 | 5.6% | 1226 | 22.5% |
| **4** | 4180 | 2723 | 65.1% | 306 | 7.3% | 230 | 5.5% | 921 | 22.0% |
| **5** | 6332 | 4183 | 66.1% | 410 | 6.5% | 394 | 6.2% | 1345 | 21.2% |
| **Total** | 27655 | 17858 | 64.6% | 1942 | 7.0% | 1600 | 5.8% | 6255 | 22.6% |

**Table 5.**
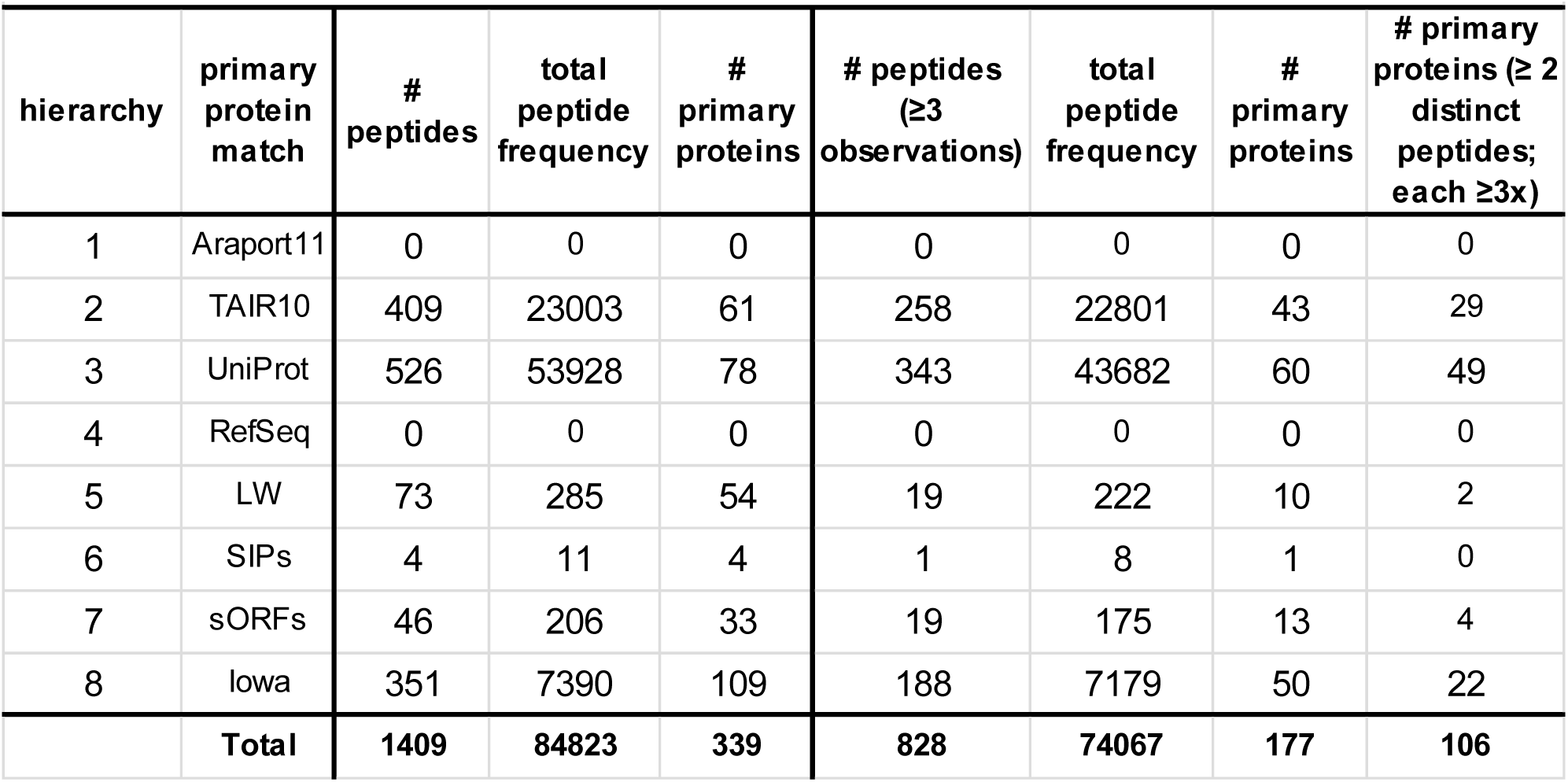
Peptide and proteins identified not identified in Araport11 but identified in one or more of the other sources.

| hierarchy | primary protein match | # peptides | total peptide frequency | # primary proteins | # peptides (≥3 observations) | total peptide frequency | # primary proteins | # primary proteins (≥ 2 distinct peptides; each ≥3x) |
| --- | --- | --- | --- | --- | --- | --- | --- | --- |
| 1 | Araport11 | 0 | 0 | 0 | 0 | 0 | 0 | 0 |
| 2 | TAIR10 | 409 | 23003 | 61 | 258 | 22801 | 43 | 29 |
| 3 | UniProt | 526 | 53928 | 78 | 343 | 43682 | 60 | 49 |
| 4 | RefSeq | 0 | 0 | 0 | 0 | 0 | 0 | 0 |
| 5 | LW | 73 | 285 | 54 | 19 | 222 | 10 | 2 |
| 6 | SIPs | 4 | 11 | 4 | 1 | 8 | 1 | 0 |
| 7 | sORFs | 46 | 206 | 33 | 19 | 175 | 13 | 4 |
| 8 | Iowa | 351 | 7390 | 109 | 188 | 7179 | 50 | 22 |
|  | **Total** | **1409** | **84823** | **339** | **828** | **74067** | **177** | **106** |

At the highest level of confidence are the canonical proteins (in both systems; Table 2A,B) and we identified 17857 canonical proteins in Araport11 (Table 4; Supplemental Dataset 2). These canonical proteins have at least two uniquely mapped non-nested peptides of at least 9 residues (Figure 6A). This is a very high standard of identification and follows the HPP guidelines (Deutsch et al., 2019). We note that if gene loci were represented by different protein isoforms (gene models), we assigned one isoform as the canonical protein and did not further count the other isoforms, unless there was a uniquely mapped peptide to the alternate protein model. Unless a higher isoform number (gene model) received stronger MS support, isoform #1 was selected. In 878 cases, the canonical protein was an isoform with a higher model number (653 for .2; 99 for .3; 25 for .4; 10 for .5; no identification of isoform .6 or higher was observed even for genes that have up to 27 isoforms!). Inspection of these genes for which a higher isoform number was the canonical form showed a range of scenarios that explain the specific identification of the alternative isoform instead of the default .1. These included extra N-terminal or C-terminal protein sequence or additional internal exon due to different splicing. Most isoforms are very similar or even identical at the protein level, and in many cases it was very hard or even impossible to distinguish between protein models based on MSMS data.

We also identified 1943 Araport11 proteins in the ‘uncertain’ category encompassing tiers 2-7 (Supplemental Dataset 2). These proteins have too few uniquely-mapping peptides of ≥ 9 aa to qualify for canonical status and may also have one or more shared peptides with other proteins. We identified 1600 Araport11 proteins assigned to the ‘redundant’ category encompassing tiers 8 and 9 (Supplemental Dataset 2). These proteins have only peptides that can also be assigned to other entries and thus these proteins are not needed to explain the observed peptide evidence. Finally, there were 6255 (6255/27655 = 22.6%) predicted proteins in Araport11, quite evenly distributed across the five nuclear chromosomes, for which we did not observe any peptides above our minimum PSM significance threshold (‘not observed’ or tier 10) (Table 4) (Supplemental Dataset 2). Some of these ‘not observed’ proteins may have low significance PSMs but these are not considered as evidence for identification for the PeptideAtlas. To better understand the nature of these unobserved proteins, we will compare the physicochemical properties and functions for these unobserved proteins and compare them with the canonical proteins in a section further below. In the remainder of the current section, we will show examples of identification of Araport11 proteins in the ‘uncertain’ category (tiers 2-8) (Figures 4B, 4C and 4D).

Within tier 2 (indistinguishable representative), we identified at a high level of confidence 27 groups of different proteins (each with unique primary sequences within the group) but for which all group members were identified by the same set of shared peptides (Figure 6A). At least some of the group members must have been detected but it is not possible to determine which ones based on the detected peptides. In most cases, members of these groups share significant sequence identity/similarity, and they often have similar types of functions. One protein was selected as the representative for each group and was placed into tier 2 and the others in tier 7. An example of this scenario is the plastid-localized family of nucleoid-interacting protein PTAC18 (AT2G32180) (selected as indistinguishable representative in tier 2), PTAC18-like (AT2G32650; selected as entry in tier 7) and AT2G27402 (tier 8 – subsumed), as shown in figure 6B. PTAC18 and PTAC18-like differ by only 2 amino acids in their protein sequences (16 kDa, 139 aa), whereas AT2G27402 is a much smaller protein (6 kDa) which high sequence identity to PTAC18. A set of overlapping and/or nested peptides matched to an N-terminal region in all three proteins, whereas as several other peptides matched to PTAC8/PTAC8-like only but they did not cover their slight differences.

Tier 3, with 309 groups of different proteins identified (each protein with unique primary sequences) is similar as tier 2, but here the situation was more complex with group members sharing one or more matched peptides and none has uniquely-mapping peptides (Figure 6A). Again, one representative member of each group was selected and assigned to tier 3 and the other group members were assigned to tier 7. 576 groups belonging to the tier 4 ‘Marginally distinguished’ were identified. Proteins in this category share several peptides with a canonical protein, but also have one uniquely mapping peptide of ≥9 residues (Figure 6A). Exploring tier 4, we noticed that in many cases the uniquely mapping peptide differed by a single aa change to a mapped peptide of the canonical protein in the same group. Consequently, this requires careful inspection of the underlaying MSMS spectra paying particular attention of the coverage by b and y ions of the key peptides.

978 proteins (tier 5 ‘Weak’) were identified that had at least one uniquely mapping peptide of ≥9 residues but that did not meet the criteria for canonical (Figure 6A). Figure 6C shows the example of AT1G27330.1 of such a tier 5 identification. This is a small Ribosome Associated Membrane Protein (RAMP4) (68 aa) with one predicted transmembrane domain in the C-terminal portion, a positive GRAVY index (0.034) and three nested or overlapping peptides, each identified multiple times across several independent PXD datasets and publications. Moreover, the N-terminal region contains eight closely spaced lysine and arginine residues which would generate very short (3-5 aa) peptides that are too small to be considered as supporting evidence by MSMS. Therefore, whereas this protein was not considered a canonical identification (tier 1) and only a tier 5 identification, this constitutes a rather solid identification. We do note that most other identified proteins in this category only have a single distinct peptide (sometimes called ‘one hit wonders’ – see (Cottingham, 2009)) and are therefore typically less reliable (even if the peptide was identified multiple times).

53 Araport11 proteins were identified in Tier 6 (Insufficient evidence) and they have one or more uniquely mapping peptide but none reach 9 residues in length (Figure 6A). We note that most MS-based studies allow peptides as short as 7 amino acids for protein identification, but shorter peptides are generally not considered. Hence the nine aa criterium applied here is relatively stringent. Figure 6D shows a tier 6 example of a steroid dehydrogenase (ATDET2/ DWARF6; AT2G38050.1) involved in the brassinolide biosynthetic pathway. It has five or six predicted transmembrane domains and a positive GRAVY index of 0.132. It was identified in two publications across some 40 different sample types with a nine aa N-terminal peptide (just downstream of a short hydrophobic region (perhaps part of the signal peptide) and one eight aa C-terminal peptide. Whereas this was not a canonical identification (since both of the peptides were only 8 aa in length), this appears to be a fairy robust identification, in particular considering that most of the protein does not yield suitable tryptic peptides for MSMS analysis. Nearly all other identifications in this tier 6 are based on a single distinct peptide are therefore potentially less reliable (‘one-hit wonders’ as in tier 5). However, several recent large-scale papers aiming to obtain a deep coverage of cellular proteomes provide experimental support (*e.g.* by MRMs or PRMs) that these so-called ‘one-hit-wonders’ can represent true identifications (Chen et al., 2014; Vandenbrouck et al., 2016). Therefore, Arabidopsis proteins identified in tiers 5 and 6 are valuable to expanding proteome coverage but require close manual scrutiny before being used as experimental support.

69 proteins (tier 7 – indistinguishable) and 1388 proteins (tier 8 – subsumed) were identified based on one or more matched peptides. However, none of these peptides were uniquely mapped and none of these proteins were selected to be the representative of a group of identified proteins (see Figure 6A). Finally, 143 proteins were assigned to tier 9; a protein in this tier has an identical protein sequence to another one (Figure 6A), and this one is effectively removed from category competition, meaning that its partner can achieve a higher status such as canonical since it is not competing with identical sequences for uniqueness mapping.

Similar as in all other plants, Arabidopsis also has a small plastid genome and mitochondrial genome. Most sources recognize 88 protein coding genes on the Arabidopsis plastid genome, with the initial sequence reported in (Sato et al., 1999), and typically 33 protein coding genes on the mitochondrial genome (Sloan et al., 2018). To our surprise (realized at the last stage of completing this first build), Araport11 (and also TAIR10) includes 122 predicted mitochondrial-encoded proteins (annotated as ATMGxxxxx). Comparison of these 122 protein sequences with the recently updated sequences (33 in total) from (Sloan et al., 2018) shows that only a subset does match. Several plastid- and many mitochondrial-encoded mRNAs undergo mRNA editing and/or trans-splicing which can affect the resulting protein sequence, thus increasing the protein search space (Takenaka et al., 2013; Germain et al., 2015; Fuchs et al., 2020; Small et al., 2020). We have reached out to members of the plant community for input and advice to obtain the most complete set of possible organelle-encoded proteins, including their unedited and edited variants. We will revisit protein accumulation, including partial editing and possible tissue specificity, of these organelle-encoded proteins in a follow-up study. In the current build a total 59 and 15 Araport11 plastid- and mitochondrial proteins respectively were identified at the highest confidence level (‘canonical’) (Table 4). For seven plastid and 80 mitochondrial predicted proteins, we did not observe any matched MSMS spectra.

### Identification and discovery of proteins not represented in Araport11

We identified 1408 peptide sequences (length at least seven amino acids; irrespective of PTM or charge state) that did not match to Araport11 protein sequences but instead matched to predicted amino acid sequences in one or more of the other sources listed in Table 1 (Supplemental Dataset 3A). The number of observations of these peptides ranged from 1 (408 peptides) to 8854. Figure 7 shows a frequency distribution for the number of peptide observations (PSMs). These peptides matched to 339 primary protein identifiers from the different sources, *i.e*. 61 in TAIR10, 78 UniProtKB, 54 LWs, 4 SIPs, 33 sORFs, 109 from Iowa (Table 5; Supplemental Datasets 3B-E). When removing peptides only observed once or twice, the number of observed protein identifiers is reduced and are: 43 in TAIR10, 60 UniProtKB, 10 LWs, 1 SIP, 13 sORFs, 50 Iowa, totaling 177 proteins. Requiring at least 2 unique peptide sequences to further reduce false discovery, the number of identified proteins was reduced to 106 (Table 5). It should be noted that that we applied a strict hierarchy to assign peptides to protein sequences from these additional sources. That is, even if a peptide was matched to a protein sequence in more than one source, the peptide was assigned to the sequence in the most highly ranked source (for ranking see Table 5). For instance, a peptide matched to a sequence in TAIR10 would not be used again to report a sequence in UniprotKB. In the next sections we explore the significance for some of these 106 proteins sequences.

**Figure 7.**
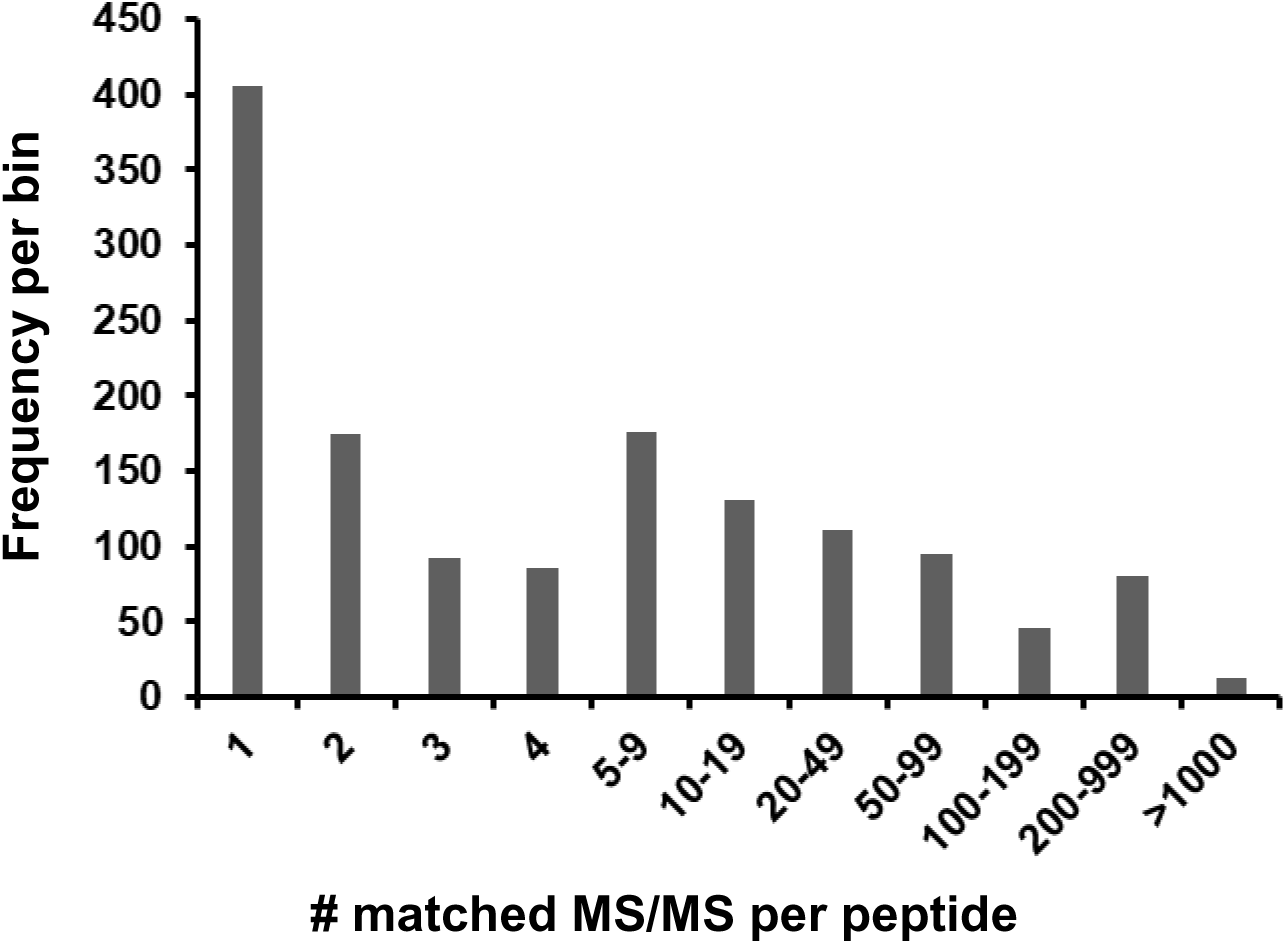
Frequency of observation for peptides not matching to Araport11 entries, but matching to other proteins sources, including TAIR10, UniProt, sORFs and other contributions.

### Proteins identified in TAIR10 and absent in Araport11

There are 32785 distinct predicted protein sequences in TAIR10 (represented by 35386 gene models) of which 1651 do not have 100% identical protein sequences in Araport11. As indicated in Table 5, we identified uniquely mapping peptides for 61 proteins in TAIR10 that could not map to Araport11, and the number of unique matched peptide sequences per protein ranged from 1 to 172 (Supplemental Dataset 3B). 43 proteins were observed with at least one distinct peptide observed all least three times by MSMS and 29 of these were observed with at least two unique peptide sequences identified at least three times (Table 5). We compared these 29 TAIR10 genes to Araport11 genes and we observed five different scenarios that explain the peptides uniquely identified for TAIR10 proteins: i) the gene was removed from Araport11 and there was no protein coding gene in this chromosomal region (5 genes – see example in Figure 8A), ii) alternative START site; in all cases the Araport11 protein was shorter (5 genes), iii) alternative STOP site; in all cases the Araport11 protein was shorter (3 genes – see example in Figure 8B), iv) mismatch within an exon (3 genes), iv) different splicing events either due to a change in the length of the exon or due to addition or removal of an exon; in all cases there was also a change in the START and/or STOP codon (11 genes – see example Figure 8C), and vi) finally in two cases the TAIR10 protein was mitochondria-encoded. Table 6 summarizes the findings for these 27 nuclear-encoded TAIR10 proteins and they should be considered for future Arabidopsis genome annotations. Figures 8A-C shows examples by comparing the TAIR10 and Araport11 chromosomal regions, assigned gene models and uniquely matched peptides for the TAIR10 entry.

**Figure 8.**
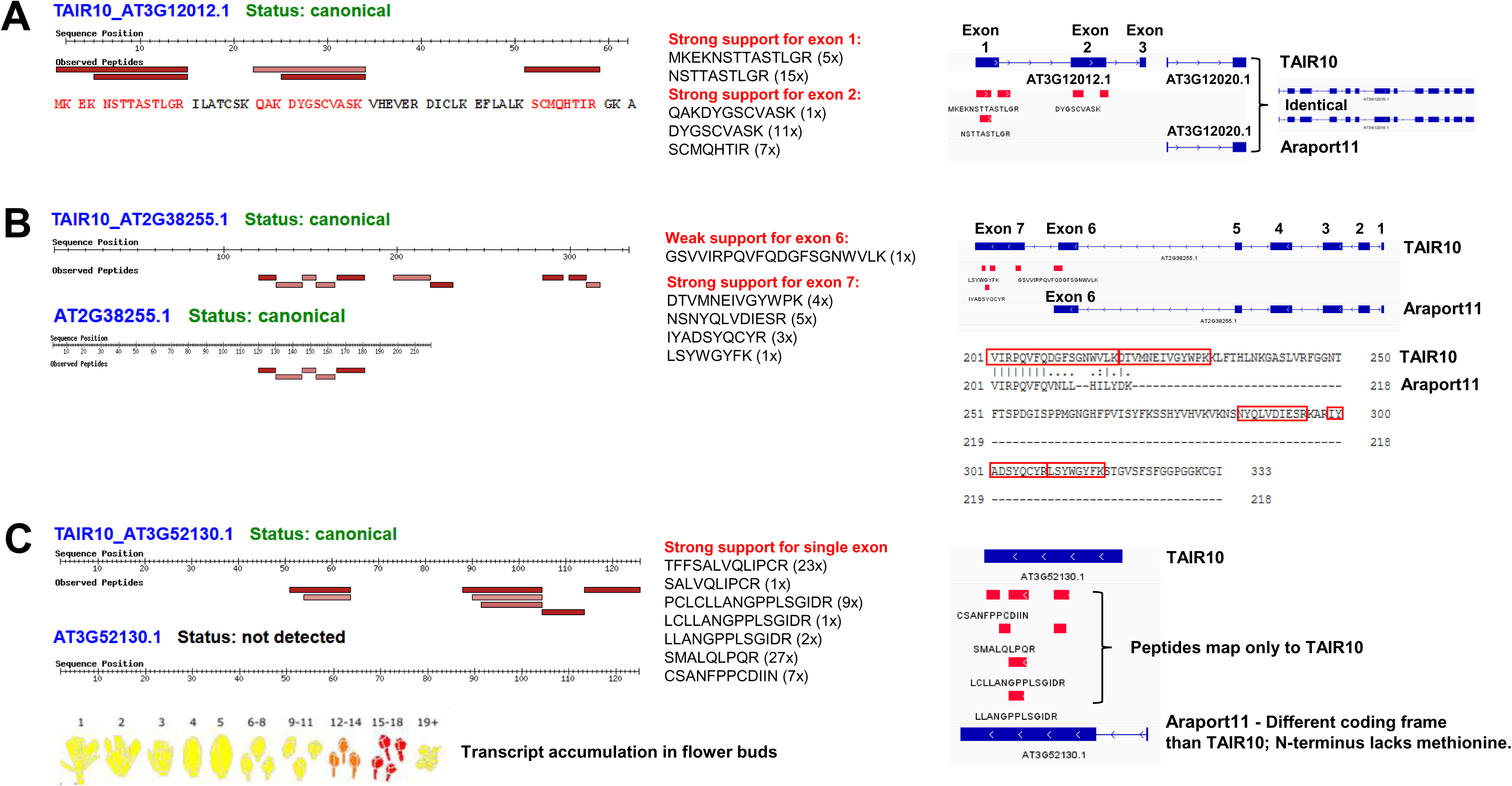
Examples of identified proteins not captured well in Araport11 but detected in TAIR10. For a complete list see Supplemental Dataset 3A,B and Table 6. **(A)** AT3G12012.1 was identified in TAIR10 but is not annotated as a protein coding gene in Araport11. The predicted protein sequence is shown with the identified residues marked up in orange (left side). The specific peptides identified by MSMS and their frequency of observation are shown (middle). The right-hand panel show the predicted gene structure with three exons in the TAIR10 annotation. This short gene is positioned within the 5’UTR of protein coding gene AT3G12010.1 and is likely an expressed uORF with unknown function. AT3G12010.1 (annotated as C18orf8; 782 aa) is identical in TAIR10 and Araport11 and was identified at the canonical level (59% sequence coverage). **(B)** Alternative protein model AT2G38255.1 (unknown protein with DUF239) with extended C-terminus in TAIR10 exhibiting multiple detected peptides not found in the shorter Araport11 entry. This was due to an alternative STOP codon combined with a change in splicing. Consequently, TAIR10 AT2G38255 has 7 exons in TAIR10 but 6 exons in Araport11; exon 7 is missing in Araport11 and exon 6 partially differs between TAIR10 and Araport11. The protein sequence alignment shows the C-terminal region of the TAIR10 (333 AA) and Araport11 (218 AA) proteins has 218 residues; the two sequence are identical until residue number 208. Five distinct peptides match to shared region of the TAIR10 and Araport11 entries – these are SQIWLENGPR, TGCYNTNCPGFVIISR, LTIYWTADGYK, GELNSIQFGWAVHPR, LYGDTLTR (see the PeptideAtlas for details). **(C)** Detection of the TAIR10 version of AT3G52130.1 (non-Type III lipid transfer protein), with no detection of the completely different sequence for AAT3G52130.1 in Araport11. This was due to alternative START and STOP codons combined with a change in splicing; the coding frames between the two genes are different. Consequently, in case of Araport11, the N-terminal residue is a lysine and not a methionine. mRNA accumulation is limited to young flower buds as displayed in BAR ePlant (yellow -> red scale reflects low to high expression values). Primary sequences for both proteins are: TAIR10_AT3G52130.1 (125 aa): MMMKAMRVGLAMTLLMTITVLTIVAAQQEGLQQPPPPPMLPEEEVGGCSRTFFSALVQLIPCR AAVAPFSPIPPTEICCSAVVTLGRPCLCLLANGPPLSGIDRSMALQLPQRCSANFPPCDIIN Araport11_AT3G52130.1 (123 aa): RSKRACNNHLHHQCCPRRKWEDAAGHFSPRWYSSYHVEQQLLLLARSHRPRYVALPLSHLV VLVFASLPMDLHSLALTAPWLFSSLRDALLISLPAISSTRKDISSFFSFLFSFTFLFNNLAA

**Table 6.** 27 nuclear-encoded TAIR10 proteins identified based on at least 2 distinct peptides supported at least three MSMS spectra, but not identified in Araport11.

| TAIR10 | # unique matched peptide sequences (includes also peptides observed only 1x or 2x) | # total frequency of observation of matched peptides | protein name | Change in Araport11 as compared to TAIR10 | Comments | # gene models in TAIR10 | # gene models in Araport11 |
| --- | --- | --- | --- | --- | --- | --- | --- |
| AT1G14070.1 | 12 | 28 | Xyloglucan fucosyltransferase | removed | 2 exons; very strong support 2nd exon | 1 | n/a |
| AT2G33430.1 | 15 | 1339 | MORF2 (also DAG protein) - editing factor | removed | very strong peptide support for 3 of the 3 exons | 1 | n/a |
| AT3G12012.1 (Figure 8A) | 5 | 39 | conserved peptide upstream ORF20 | removed | 3 exons. strong support for 2 exons | 1 | 0 |
| AT4G18120.1 | 22 | 202 | MEI2-like 2 | removed | very strong peptide support for 7 of the 10 exons. How are model .1 and .2 different (Tair10) | 2 | n/a |
| AT5G25752.1 | 7 | 235 | rhomboid protease 11/9 (RBL11/9) | removed | 7 exons (Tair10). 5 peptides for first exon. One peptide for last exon | 1 | n/a |
| AT1G16780.1 | 3 | 23 | Inorganic H pyrophosphatase | ATG shifted. Araport11 is shorter | 15 (Tair10) or 14 exons (Ara11). All Araport models start later | 1 | 3 |
| AT5G18280.2 | 3 | 478 | apyrase 1 | ATG shifted. Araport11 is shorter | 9 exons in model 1 and 11 in model 2 for Tair10. 9 exons in Araport11. one extra exon in model .2 supported with one short peptide; 2 peptides for first exon. | 2 | 1 |
| AT5G52640.1 | 4 | 51 | heat shock protein (HSP81-1/HSP83) | ATG shifted. Araport11 is shorter | 4 exons in both Tair10 and araport11; strong support for 1st exon | 1 | 1 |
| AT5G63980.1 | 4 | 54 | SAL1 (FIERY1), 3(2),5-bisphosphate nucleotidase | ATG shifted. Araport11 is shorter | 7 exons in both Tair10 and Araport11; 4 peptides for the 1st exon | 1 | 1 |
| AT5G18280.1 | 3 | 478 | apyrase 2 | ATG. Araport11 is shorter | 9 exons in model 1 and 11 in model 2 for Tair10. 9 exons in Araport11. one extra exon in model .2 supported with one short peptide; 2 peptides for first exon. | 2 | 1 |
| AT1G03495.1 | 4 | 26 | HXXXD-type acyl-transferase | STOP. Araport11 is shorter | one big exon - clear case | 1 | 1 |
| AT1G79920.1 | 26 | 2006 | Heat shock protein 70 (Hsp 70) | STOP. Araport11 is shorter | 9 exons in Tair10; 8 exons in Araport11 | 2 | 4 |
| AT4G23000.1 | 5 | 16 | Calcieneurin-like metallo-phosphoesterase | STOP. Araport11 is shorter | 15 exons in Tair10, 14 exons in Araport11. several peptides for both extra exons | 1 | 2 |
| AT5G40780.2 | 3 | 21 | lysine histidine transporter 1 | splicing | 8 exons in Tair10; 8 exons for model 1, but 7 for models 2 and 3 in Araport11. | 2 | 3 |
| AT3G57180.1 | 3 | 28 | BPG2- homologue of Yqeh-GTPase | mismatch - in one exon | 3 overlapping long peptides; sequencing difference? | 1 | 1 |
| AT4G16150.1 | 3 | 18 | calmodulin-binding transcription activator 5 | mismatch - in one exon; likely related to a gene duplication AT3G16940 & AT4G16150 | 13 exons in both Tair10 and Araport11. sequencing mismatch in 11th exon | 1 | 1 |
| AT1G52827.1 | 3 | 191 | cadmium tolerance 1 | ATG+STOP. Big change | 3 nested peptides for exon 1; no peptides for exon 2 | 1 | 1 |
| AT3G52130.1 (Figure 8C) | 7 | 70 | Bifunctional inhibitor/lipid-transfer protein/seed storage 2S albumin | ATG+STOP+splicing | one exon in Tair10; 2 exons in Araport - but first new one very short | 1 | 1 |
| AT4G16770.1 | 3 | 66 | 2-oxoglutarate (2OG) and Fe(II)-dependent oxygenase | ATG+STOP+splicing. Araport11 is shorter | 11 exons in Tair10 and 8 in Araport11. 2 peptides for extra C-terminal exon; splice difference for exon 7 | 1 | 1 |
| AT5G39570.1 | 172 | 14860 | transmembrane protein | ATG+STOP+splicing. Araport11 is shorter | 2 exons in Tair10; 2 or 3 exons in Araport11. | 1 | 2 |
| AT4G21326.1 | 3 | 12 | subtilase SBT12 | ATG+splicing. Araport 11 is shorter | 8 exons in Tair10, 7 exons in Araport11 | 1 | 1 |
| AT1G23580.1 | 10 | 49 | transmembrane protein with DUF220 | ATG+splicing. Araport11 is shorter | 4 exons in Tair10; 2 exons in Araport11 | 1 | 1 |
| AT2G38255.1 (Figure 8B) | 5 | 14 | hypothetical protein (DUF239) | STOP+splicing | 7 exons (Tair10); 6 exons in (Araport11). Strong support for extra C-term exon | 1 | 1 |
| AT1G17060.1 | 7 | 33 | cytochrome p450 72c1 | STOP+splicing | 6 exons (TAIR10) or 4 exons (Araport11); Araport 11 - antisense gene - overlapping protein coding - AT1G7065) | 1 | 1 |
| AT4G16144.1 | 5 | 46 | JAB1/Mov34/MPN/PAD-1 domain protein | STOP+splicing. Araport11 is shorter | 13 exons in Tair10 and 13 in Araport11. 4 peptides in extra C-term exon, and 1 for splice junction | 1 | 1 |
| AT4G18260.1 | 11 | 103 | Cytochrome b561/ferric reductase | STOP+splicing. Araport11 is shorter | 7 exons in Tair10, 4 exons in Araport11. All three extra C-terminal exons are supported with peptides | 1 | 1 |
| AT5G02370.1 | 2 | 11 | kinesin motor protein-related | STOP+splicing. Araport11 is shorter | 1 exons in Tair10 but 9 in Araport11; 2 nested peptides for the last exon | 1 | 1 |

TAIR10 AT3G12012 is represented by one gene model and it has three exons (Figure 8A). The MSMS data provide very strong support for exons 1 and 2, by two and three peptides respectively, but not for the very short exon 3 which encodes for just three residues (GKA) immediately downstream of a lysine residue. This 3rd and C-terminal exon can only be observed if there were two missed cleavage in the C-terminal region resulting in the peptide SCMQHTIRGKA. This annotated gene is not present in the Araport11 genome annotation and the description in the TAIR database states that the gene is obsolete but previously it was assigned as a conserved upstream opening reading frame (uORF) named CPuORF20 in the 5’UTR of the protein coding gene AT3G12010 (annotated as C18orf8 in TAIR - Araport11) (Figure 8A). We identified AT3G12010.1 as a canonical protein with very high sequence coverage and the TAIR10 and Araport11 protein sequences are identical. It appears that AT3G12010 is indeed a short uORF that is expressed as a stable small protein (6.7 kDa) with unknown function.

TAIR10 AT2G38255 (unknown protein with DUF239) has 7 exons in TAIR10 but 6 exons in Araport11; exon 7 is missing in Araport11 and exon 6 partially differs between TAIR10 and Araport11 due to a splicing difference and presence of a STOP codon in Araport11 (Figure 8B). As illustrated in the alignment, the TAIR10 protein has 333 residues and the Araport11 protein has 218 residues; the two sequence are identical until residue number 208. The MSMS data strongly support exon 7 with four distinct peptide sequences and the alternate exon 6 from TAIR10 with one MSMS spectrum. Five distinct peptides match to the N-terminal portion which is identical across TAIR10 and Araport11 (see legend Figure 8B). mRNA expression levels appear to be very low (no values are reported in *e.g.* BAR ePlant (http://bar.utoronto.ca/) or the ATTED co-expression database (https://atted.jp/), which perhaps explains the incomplete annotation of its predicted protein sequence.

AT3G52130.1 was detected in TAIR10 with seven distinct peptides and a total of 70 MSMS spectra, but not at all in Araport11 (Figure 8C). The predicted proteins sequences in TAIR10 and Araport11 are completely different due alternative START and STOP codons combined with a change in splicing; the coding frames between the two genes are different. Consequently, in case of Araport11, the N-terminal residue is a lysine and not a methionine. Both primary protein sequences are listed in the legend (Figure 8C). A study in 2013 demonstrated that AT3G52130 is a non-Type III lipid transfer protein with transcripts nearly exclusively present in the inflorescence (flower bud stage 9) (Huang et al., 2013) as also shown in BAR ePlant (Figure 8C). The TAIR10 gene assignment appears correct.

### Peptides matching to UniProtKB sequences

526 peptides did not match to Araport11 nor TAIR10 protein coding sequences but matched to 78 UniProtKB identifiers (Supplemental Dataset 3C; Table 5). Considering only peptides identified at least three times, reduced this to 343 peptides matching to 60 UniprotKB identifiers. Increasing the stringency further by requiring at least two distinct peptides (each observed at least three times), reduced the number of identified UniProtKB sequences to 49. To investigate the nature and significance of these UniProtKB identifications, we blasted all UniprotKB sequences to Araport11 protein coding genes (pBlast), to pseudogenes (tBlastn) and to the genomic sequence (tBlastn) (Supplemental Dataset 3D). We then evaluated the 49 UniprotKB identifiers that passed the stringent criteria.

Four UniprotKB ids mapped each to a different plastid-encoded protein. In three cases (ACCD, cytb559-beta and ClpP1), the unique peptides matched only to UniprotKB and not Araport11 and always included an RNA edited site because the Araport11 sequences did not consider the resulting amino acid change whereas the UniprotKB sequences were corrected for the edited site. The fourth case was for YCF3 and was due to a miss-assigned N-terminus in Araport11; instead of the correct 168 aa protein, the Araport11 sequence was only 126 aa long (the first 42 aa N-terminal residues were missing). 25 UniProtKB identifications mapped to 19 Araport11 mitochondrial-encoded proteins with 5 Araport11 proteins matching to two different UniProtKB entries. In most cases this is due to unedited forms in Araport11 and edited form in UniProtKB. As mentioned earlier, we will further investigate the coverage of the plastid- and mitochondrial-encoded proteome in a follow-up study. Twenty UniProtKB identifications mapped each to a nuclear-encoded protein in Araport11. In a handful of cases these best mapped to a pseudogene in Araport11 – but these are likely to be actual protein coding genes (*e.g.* AT3G0875, AT4G18120, AT4G13900, AT4G204033, AT4G14610). In other cases there was one more mismatch between the UniProtKB entry and the Araport11 protein, likely related to sequence annotation. Figure 9 shows three examples of UnipProtKB identifications that were selected because the UniProtKB protein sequence matched with relatively high significance to a predicted pseudogene in Araport11 (based on TBlastN). The first example is Q9LHK4 which encodes for a large 137 kDa protein (1241 aa) (Figure 9A). TBlastN identified Araport11 AT3G30875 as a strong match. AT3G30875 is annotated as a pseudogene but the TAIR website notes that this is probably not a pseudogene based on evidence for transcription (RNA-seq) and translation (Ribo-seq) described in (Hsu et al., 2016). We identified six peptides uniquely mapping to Q9LHK4, of which two were observed 3 times, and the others only once or twice. An additional three matched peptides were shared with other proteins in protein group AT4G17140 which is a protein with repeating coiled regions of VPS13. The second example is Q9SVV9 encoding for an 85 kDa protein (759 aa) and is identical to TAIR10 AT2G18120 with the exception of two gaps in the aa sequence alignment. This protein (AML3) is a member of the MEI2-like gene family and is annotated as a pseudogene in Araport11. In situ hybridization detected expression during early embryo development but not in vegetative and floral apices (Kaur et al., 2006). The third example is P0CC32 which encodes a 57 kDa protein (512 aa) and maps to the pseudogene AT2G04033 with similarity to the defensin-like (DEFL) family. However, careful inspection of the results for P0CC32 in PeptideAtlas shows that the UniProtKB sequence is nearly identical to a much smaller (42 kDa) chloroplast-encoded NDHB/NDH1 protein (ATCG00980) with the exception of an N-terminal region of 123 amino. P0CC32 was identified with two unique peptides that did not match to AT2G04033 because this protein is RNA edited and these two peptides contain one or two editing sites resulting in amino acid changes. As mentioned earlier, the Araport11 sequences are the unedited form whereas UniProtKB does incorporate these edits. Three additional peptides were identified for P0CC32 but these were all shared with AT2G0433.

**Figure 9.**
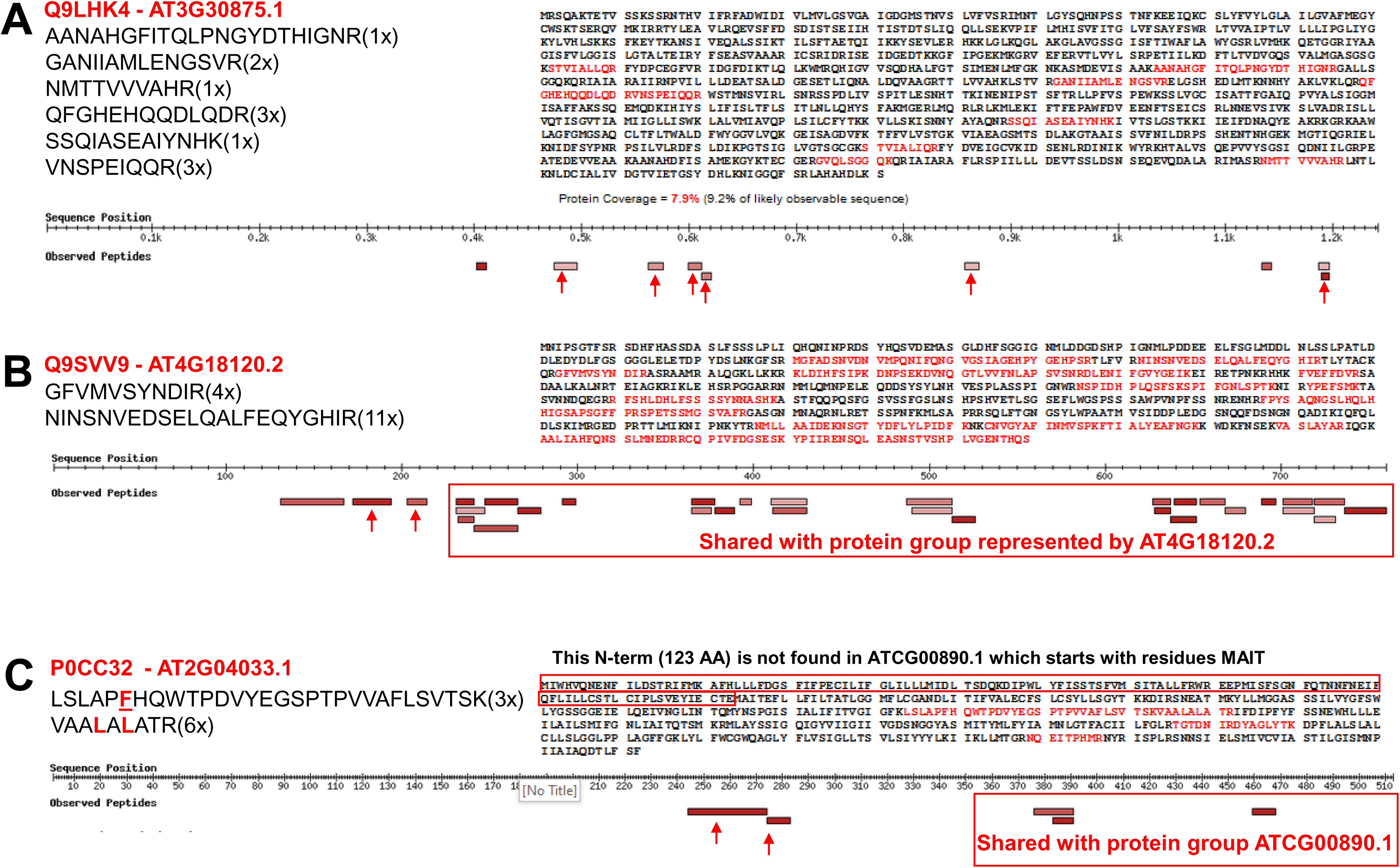
Examples of UniProtKB identifications selected from the 49 identifications based on at least two distinct peptides (not matched to predicted Araport11 proteins) that were observed at least three times. For a complete listing see Supplemental Dataset 3D. The examples were selected because the UniProtKB protein sequence matched with relative high significance to a predicted pseudogene in Araport11 (based on TBlastN). **(A)** Q9LHK4 encodes for a large 137 kDa protein (1241 aa). TBlastN identified Araport11 AT3G30875 as a strong match (1215 aa alignment length and E = 0). AT3G30875 is annotated as a pseudogene (nine exons are shown in Jbrowse) but the TAIR website notes that this is probably not a pseudogene based on evidence for transcription (RNA-seq) and translation (Ribo-seq) described in (Hsu et al., 2016). We identified six peptides (marked with red arrows) uniquely mapping to Q9LHK4 (sequences are shown), of which two were observed 3 times, and the others only once or twice. An additional three matched peptides were shared with other proteins in protein group represented by AT4G17140.3, which is a protein with repeating coiled regions of VPS13. **(B)** Q9SVV9 encodes for an 85 kDa protein (759 aa) and is identical to TAIR10 AT4G18120.2, with the exception of two gaps in the aa sequence alignment. This protein (AML3) is a member of the MEI2-like gene family and is annotated as a pseudogene in Araporti1. *In situ* hybridization detected expression during early embryo development but not in vegetative and floral apices (Kaur et al., 2006). Two peptides uniquely map to Q9SVV9 as indicated by the red arrows. The other matched peptides are shared with the protein group represented by AT4G18120.2 as indicated. **(C)** P0CC32 encodes a 57 kDa protein (512 aa) and maps to the pseudogene AT2G04033.1 with similarity to the defensin-like (DEFL) family. However, careful inspection of the results for P0CC32 in PeptideAtlas shows that the Uniprot sequences is nearly identical to much smaller (42 kDa) chloroplast-encoded NDHB/NDH1 protein (ATCG00980.1) with the exception of an N-terminal region of 123 amino. P0CC32 was identified with two unique peptides that did not match to AT2G04033.1 because this protein is RNA edited and these two peptides contain one or two editing sites resulting into aa changes. As mentioned in the Results and Discussion section, the Araport11 sequences are in the unedited form whereas UniProtKB does incorporate these edits. Three additional peptides were identified for P0CC32 but these were all shared with AT2G0433.1 as indicated.

### Discovery of small Open Reading Frames (sORFs)

Transcriptomics, including using Ribo-seq, combined with a range of *in silico* prediction and analysis tools have predicted large numbers of small open reading frames (sORFs) in the Arabidopsis genome that could result in accumulation of small proteins or peptides (Hanada et al., 2007; Hsu et al., 2016; Hazarika et al., 2017; Hsu and Benfey, 2018; Takahashi et al., 2019; Kage et al., 2020). These sORFs have been found in intergenic regions, introns, embedded within non-coding RNAs (ncRNAs), directly upstream of coding sequences in the 5’ untranslated regions (uORFs), C-terminally encoded peptides (CEPs) (Roberts et al., 2013), or on the anti-sense strand and some are induced by (a)biotic stresses, sometimes assigned stress induced peptides (SIPs) (Hazarika et al., 2017; Qi et al., 2020; Takahashi et al., 2020). Recent mass spectrometry studies searched these ORF collections for Arabidopsis and identified matching peptides for a relative low percentage of predicted proteins or peptides (Zhang et al., 2019; Mergner et al., 2020; Wang et al., 2020). The assignments within this collection are low weight proteins (LWs), SIPs and sORFs. As indicated in Table 5 and Supplemental Dataset 3D, we identified 54, 4 and 33 LWs, SIPs and sORFs based on 74, 4 and 46 peptides, respectively. When considering only peptides observed at least three times, this was reduced to 10, 1 and 13, LWs, SIPs and sORFs respectively. Upon increasing the stringency by requiring two distinct peptides, each observed at least three times, only four sORFs and 2 LWs remained and we investigated these six most stringent hits. Figure 10 shows the identification of these elements by displaying screenshots for their identifications in PeptideAtlas showing how the peptides map to the predicted protein sequence, and how they map to the Arabidopsis genome sequence.

**Figure 10.**
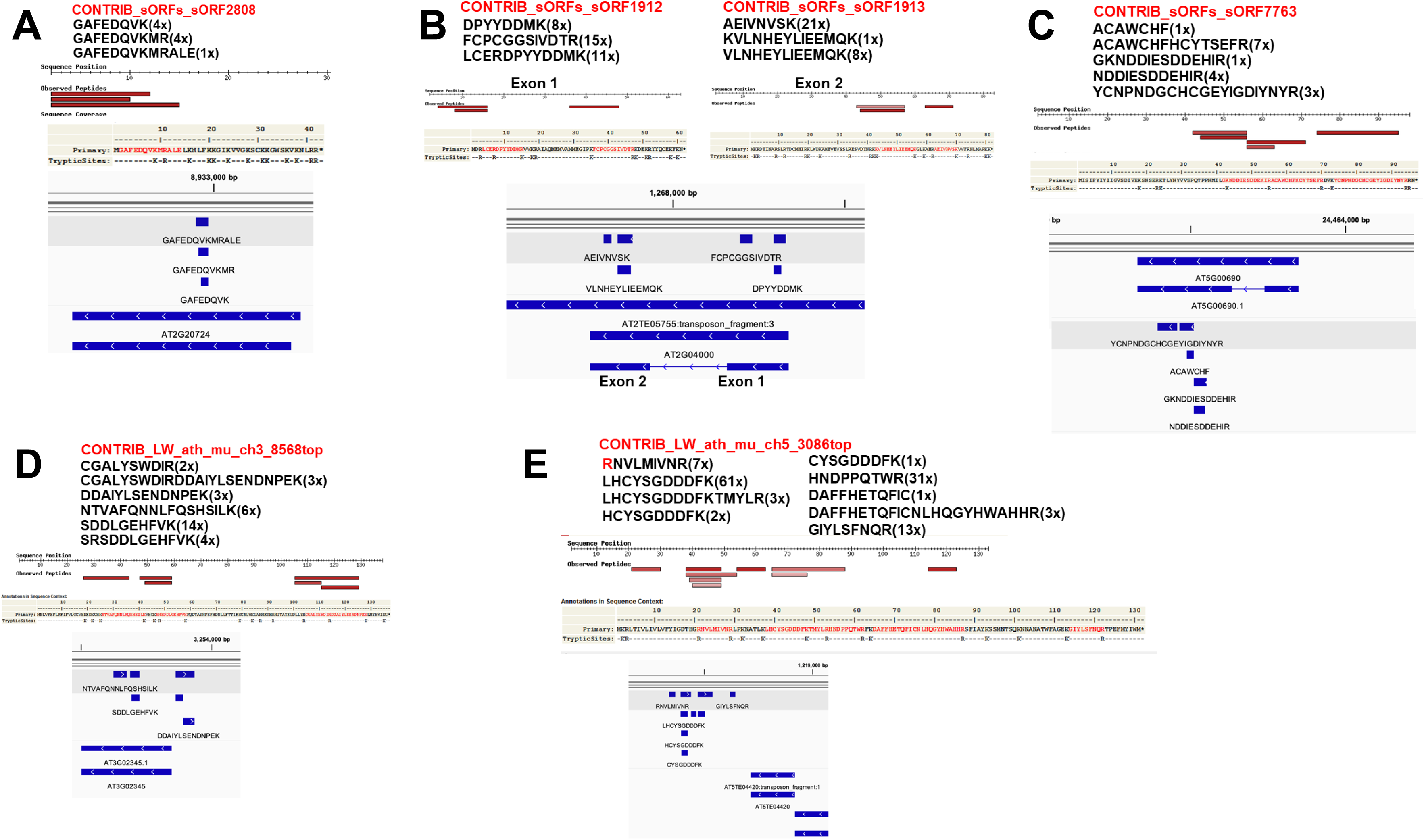
Examples of identification of sORFs and one LW in the PeptideAtlas build. These six examples represent the identifications that pass the stringent criterium of having at least two matched distinct peptides that are each identified at by at three times (for more information, see Supplemental Dataset 3D). **(A)** CONTRIB_sORFs_sORF2808 encodes for a 4.9 kDa peptide (42 aa) and was identified with three distinct and nested peptides that map to a small portion of the pseudogene AT2G20724 in Araport11. The identified peptides were all identified from dimethyl labeling (modifying both N-terminal free amines and the lysine side chain) and enrichment studies using TAILS or COFRADIC) as indicated. The 4.9 kDa predicted protein has a very high number of lysine and arginine residues (total 13) and upon tryptic digestion would result in only a single peptide of 7 (N-terminal methionine removed) or 8 amino acids. Trypsin cannot cleave the peptidyl bond of dimethylated lysine residues which enhances the chance to observed peptide GAAFEDQVKMR and GAAFEDQVKMRALE. All four PSMs of GAFEDQVK and one of the four PSMs of GAFEDQVKMR are dimethylated and the other three are iTRplex labeled. The single PSM of GAFEDQVKMRALE is iTRplex labeled. The results suggest that AT2G20724 is not a pseudogene but rather a protein-coding gene. **(B)** Both CONTRIB_sORFs_sORF1912 and CONTRIB_sORFs_sORF1913 mapped to the same transposon AT2TE05755/AT2G04000 in Araport11. AT2G0400 has two exons and the two ORFs each represent one exon. The transposon belongs to the VANDAL21 family and superfamily DNA/MuDR and its preferred substrate for integration of VANDAL21 is the euchromatin. VANDAL21 targets mainly promotors and 5’UTR of broadly active genes which are enriched in histone marks H3K4me3 and H3K36me3 (Quesneville, 2020). **(C)** CONTRIB_sORFs_sORF7763 encodes for a 11 kDa protein (96 aa) and was identified with five peptides that all map to exon 2 of AT5G00690 in Araport11. However, Araport11 has not assigned this as a protein-coding gene but as a ‘novel transcribed region’. The results suggest that AT5G00690 should be annotated as a protein coding gene. **(D)** CONTRIB_LW_ath_mu_ch3_8568top encodes for a 16 kDa protein (136 aa) and was identified with six peptides, three of which mapped to AT3G02345 which is annotated as a long-non-coding RNA in Araport11. Most PSMs were identified in seeds and a few others in embryos or siliques (see PeptideAtlas). BlastP with the 136 aa sequence against Araport11 found that the closest match was AT2G23148 but with very poor E-value (0.003). BlastP against all nr proteins identified ARALYDRAFT_897225 in the lyrate ecotype as the closest match (98/117 identities for the region 20-80 aa) (1E-72). The significance of the small protein remains to be determined. **(E)** CONTRIB_LW_ath_mu_ch5_3086top encodes for a 15.6 kDa protein (131 aa) and was identified with nine peptides, none of which map to an annotated genome element in Araport11. However, BlastP against all nr proteins identified AT5G03740 in ecotype Landsberg as the perfect match. The peptides were identified in samples from flowers and flower parts (petals, pollen, sepals, stamen) as well as siliques in two studies (Zhang et al., 2019; Mergner et al., 2020), even if these studies describe that samples are from ecotype Col-0.

CONTRIB_sORFs_sORF2808 encodes for a 4.9 kDa protein (42 aa) and was identified with three distinct and nested peptides that map to a small portion of the pseudogene AT2G20724 in Araport11 (Figure 10A). The observed peptides were all identified from dimethyl labeling (modifying both N-terminal free amines and the lysine side chain) and enrichment studies using TAILS or COFRADIC) as indicated. The 4.9 kDa predicted protein has a very high number of lysine and arginine residues (total 13) and upon tryptic digestion would result in only a single peptide of 7 (N-terminal methionine removed) or 8 amino acids. Trypsin cannot cleave the peptidyl bond of dimethylated lysine residues which enhances the chance to observe peptide GAAFEDQVKMR and GAAFEDQVKMRALE. The results suggest that AT2G20724 is not a pseudogene but rather a protein-coding gene.

Both CONTRIB_sORFs_sORF1912 and CONTRIB_sORFs_sORF1913 mapped to the same transposon AT2TE05755/AT2G04000 in Araport11 (Figure 10B). AT2G0400 has two exons and the two ORFs each represent one exon. The transposon belongs to the VANDAL21 family and superfamily DNA/MuDR and its preferred substrate for integration of VANDAL21 is the euchromatin. VANDAL21 targets mainly promotors and 5’UTR of broadly active genes which are enriched in histone marks H3K4me3 and H3K36me3 (Quesneville, 2020).

CONTRIB_sORFs_sORF7763 encodes for a 11 kDa protein (96 aa) and was identified with five peptides that all map to exon 2 of AT5G00690 in Araport11 (Figure 10C). However, Araport11 has not assigned this as a protein-coding gene but as a ‘novel transcribed region’. The results suggest that AT5G00690 should be annotated as a protein coding gene.

CONTRIB_LW_ath_mu_ch3_8568top encodes for a 16 kDa protein (136 aa) and was identified with six peptides, three of which mapped to AT3G02345 which is annotated as a long-non-coding RNA in Araport11 (Figure 10D) Most PSMs were identified in seeds and a few others in embryos or siliques (see PeptideAtlas). BlastP with the 136 aa sequence against Araport11 found that the closest match was AT2G23148 but with very poor E-value (0.003). BlastP against all non-redundant proteins identified ARALYDRAFT_897225 in the lyrate ecotype as the closest match (98/117 identities for the region 20-80 aa) (1E-72). The significance of this small protein remains to be determined.

CONTRIB_LW_ath_mu_ch5_3086top encodes for a 15.6 kDa protein (131 aa), was identified with nine peptides, and maps to a region of chromosome five without annotated features in Araport11 (Figure 10E). In fact, they map downstream of transposon AT5TE04420. BlastP against all nr proteins identified AT5G03740 in ecotype Landsberg as the perfect match. The peptides were identified in samples from flowers and flower parts (petals, pollen, sepals, stamen) as well as siliques in two studies (Zhang et al., 2019; Mergner et al., 2020), even if these studies describe that samples are from ecotype Col-0. AT5G03740 in Araport11 or TAIR10 is a predicted protein coding gene but with a predicted protein sequence that is unrelated to AT5G03740 in Landsberg.

### Relative abundance, physicochemical properties, subcellular locations of the canonical and unobserved Araport11 proteome

As described above and in Table 5 about 65% of all predicted Araport11 proteins (counting one isoform per gene) were identified as canonical proteins. To better understand this canonical proteome, we analyzed the physicochemical properties and subcellular localizations of these canonical proteins and compared that to the complete predicted Araport11 proteome (selecting one protein isoform per gene) as well as the unobserved (‘dark’) proteome.

#### Physicochemical properties

We calculated molecular weight (kDa), overall hydrophobicity based on the GRAVY index (positive and negative values are hydrophobic and hydrophilic, respectively), and isoelectric point (pI) for all full-length predicted, canonical, and unobserved in Araport11. The distribution of calculated properties for each group are displayed as histograms and violin plots, whereas mean, median and mode values for molecular weight and GRAVY are also shown (Figure 11A). The canonical proteome had a higher mean and median molecular weight than the total proteome shifted by 7-9 kDa whereas the mode dramatically increased by 22.5 kDa. In contrast, the unobserved proteome was strongly skewed towards proteins of lower molecular weight as reflected by much lower values for mean, median and mode (Figure 11A). The canonical proteome showed a narrower distribution for GRAVY index values, lacking proteins with GRAVY > 1 (very hydrophobic proteins) but also lacking the most hydrophilic proteins, such as a family of very basic and small ribosomal L41 homologs (AT1G56045, AT2G40205, AT3G08520, AT3G11120, AT3G56020) as well as a very small and acidic predicted replication factor (AT5G03710). The unobserved proteins with high GRAVY values are mostly low molecular weight proteins (<10 kDa) with 1 or 2 predicted transmembrane domains. Many of these low mass unobserved proteins have no known function, but a subset are well-known small thylakoid integral membrane proteins of photosystem II (*e.g.* psbZ, psbM, psbK, psbJ). The pI plots show a bimodal distribution similar with relatively few proteins with a pI around 7.5, similarly as is generally observed for many other cellular proteomes (Schwartz et al., 2001; Kiraga et al., 2007). The general explanation for this modality is that proteins are generally least soluble near their isoelectric points and that the physiological pH within cells is typically around ∼7 to ∼7.5; hence proteins tend to be more soluble at acidic or basic pH values. The canonical proteome has a similar pI distribution as the total predicted proteome, but somewhat enriched for low pI proteins, whereas the unobserved proteome has a broader distribution for the pI distribution (Figure 11A). We conclude that pI per se is not a strong predictor for protein discovery by MSMS.

**Figure 11.**
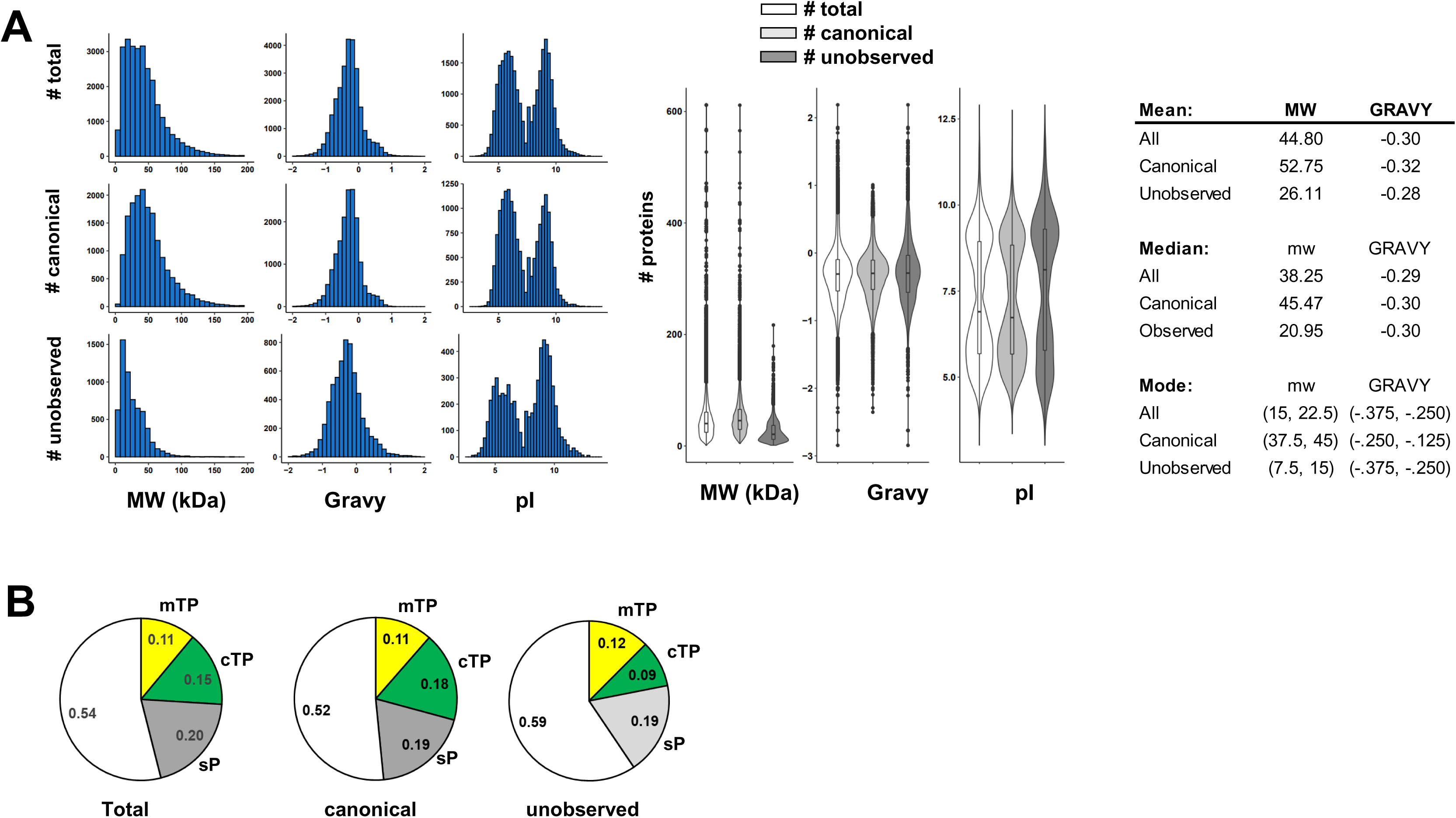
Distribution of physicochemical properties and subcellular location of the total predicted (27655 proteins, canonical (17857 proteins) and unobserved (6255 proteins) Araport11 proteome. **(A)** Frequency distributions for size, gravy, and pI for the three proteomes shown as histograms and violin plots. The table shows mean, median and mode (min-max bin value) of molecular weight and GRAVY for the three proteomes. **(B)** Distribution of subcellular localizations of nuclear-encoded proteins in the three proteomes based on predicted sP (secreted - grey), cTP (plastid - green) and mTP (mitochondria - yellow).

**Figure 12.**
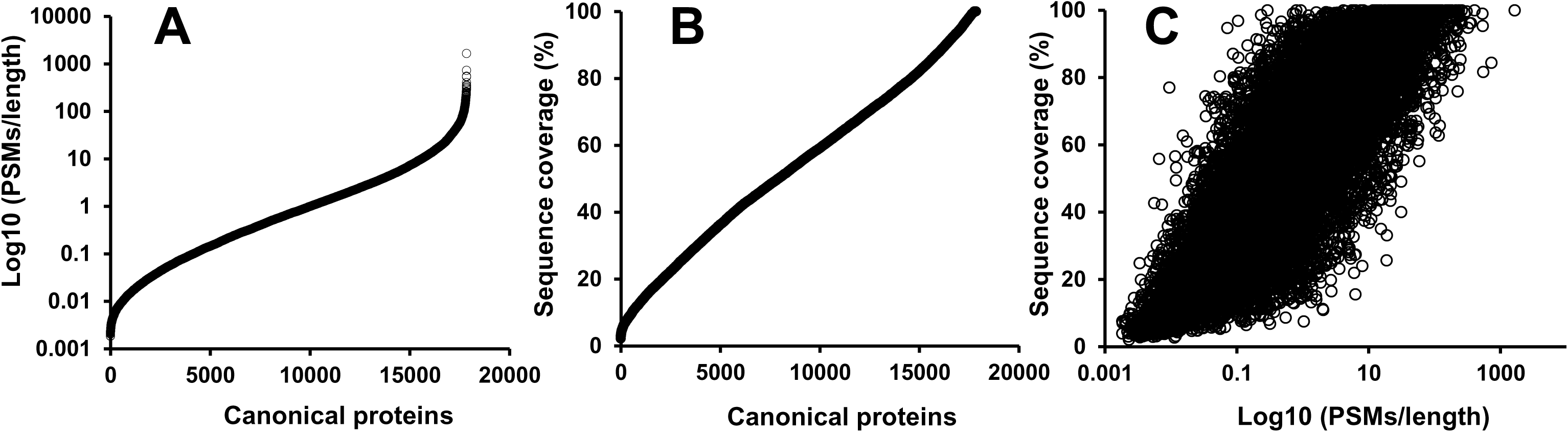
Relative abundance of the canonical proteins in Araport11 across the peptide atlas build. **(A)** Relative abundance for the canonical proteins based on the number of apportioned PSMs normalized to the length of the protein (based on number of aa). **(B)** Relative protein sequence coverage for canonical protein based on sequence coverage, *i.e.* the % of residues of the primary sequence that are part of the matched peptides (%). **(C)** Correlation between relative protein abundance (log10 (PSMs/length) and sequence coverage (%).

#### Subcellular localization

The PXDs analyzed for this first build include samples from a wide variety of plant parts and several subcellular locations (Figure 3). To get an impression of proteome coverage across subcellular localizations for the total, canonical and unobserved proteomes, we compared the predicted subcellular localizations for secreted proteins (ER, Golgi, PM, cell wall and vacuole) based on sP, plastids based on the cTP, and mitochondria based on the mTP using the well-documented localization predictor TargetP. Respectively 20%, 15% or 11% of all predicted proteins have a predicted sP, cTP or mTP. The canonical proteome was somewhat enriched for chloroplast proteins (18% cTP), whereas the unobserved proteome was strongly underrepresented in proteins with predicted cTP (9% cTP) (Figure 10C). This a rough estimate (given the uncertainties of predictions and alternative sorting mechanisms) but nevertheless suggests that the plastid proteome is relatively well covered at the canonical level.

Protein abundance determination by MS is challenging because this greatly depends on the physicochemical properties of the peptides and number of suitable peptides for a given protein (Ankney et al., 2018; Calderon-Celis et al., 2018). Accurate and absolute protein quantification is however possible, in particular when including ‘spike-in’ proteotypic peptides generated by chemical synthesis (AQUA) or through expression in *E. coli* (QConcat) (Ankney et al., 2018; Calderon-Celis et al., 2018). However, these spike-in experiments are costly and are typically done at a small scale, targeting just dozens of proteins. In the context of this PeptideAtlas, we determined relative abundance for the canonical proteins based on the number of PSMs normalized to the length of the protein (as # of aa). Furthermore, we refined that abundance by calculating the apportioned PSMs, which is the summation of the uniquely mapping PSMs and a portion of the shared PSMs based on the ratio between uniquely mapping peptides of the canonical protein and protein(s) with which the peptides were shared. The apportioned PSMs ranged from 2 to 785178, and when normalized to protein length (aa), ranged from 0.0018 to 1639, which is a dynamic range of nearly 6 orders of magnitude (Figure 11A). The five proteins with the highest relative abundance were large and small subunits of RUBISCO and CF1β of the thylakoid ATP synthase (ATCG00490, AT1G67090, AT5G38410, AT2G39730 and ATCG00480). As a complementary measure of relative abundance, we calculated the relative protein sequence coverage for canonical proteins, *i.e.* the % of residues of the primary sequence that are part of the matched peptides (%) (Figure 11B). Sequence coverage ranged from to 2% to 100%. The correlation between relative abundance and sequence coverage is positive but poor, which is expected given sequence coverage depends on suitable tryptic peptides for MSMS analysis and because many proteins accumulate without their cleavable signal peptide. Because this PeptideAtlas is built on a wide range of samples, and some tissues, subcellular fractions, or proteins are likely over- or under-sampled, and these relative abundances only provide a rough abundance estimate. However, the abundance estimate is nevertheless a useful attribute when investigating protein function.

### Post-translation modifications (PTMs)

Plant proteins undergo various physiological (*in vivo*) post-translational modifications which can often best be detected using specific enrichment methods, *e.g.* phosphorylation, acetylation, ubiquitination, SUMOylation, and cysteine (redox) modifications (Friso and van Wijk, 2015; Augustine and Vierstra, 2018; Vu et al., 2018; Sandalio et al., 2019; Moller et al., 2020). As discussed earlier, we selected PXDs that included specific peptide enrichment for phosphorylation and lysine acetylation, as well as processing events at the N-termini of proteins using various N-terminomics techniques (TAILS, COFRADIC and ChaFRADIC) in most cases combined with protein or peptide dimethylation (with/without stable isotope) to label N-terminal α-amines as well as ε-amines on the side chain of lysine (Figure 3C; Table 3; Supplemental Dataset 1). N-terminal acetylation is a very common PTM in the cytosol mostly occurring co-translationally by ribosome-associated N-terminal acetyltransferases (NATs) (Linster and Wirtz, 2018). In addition, a large portion of chloroplast-localized nuclear-as well as plastid-encoded proteins also undergo N-terminal acetylation (Zybailov et al., 2008; Rowland et al., 2015; Giglione and Meinnel, 2020). N-terminal acetylation can affect protein stability, localization and protein interactions. Lysine acetylation plays critical roles in regulating gene expression through modification of histones in the nucleus and as well as other proteins involved in a wide range of function, located across different subcellular compartments, including the cytosol, mitochondria and plastids (Hartl et al., 2017; Hosp et al., 2017; Fussl et al., 2018; Bolter et al., 2020). Phosphorylation is the most studied PTMs in Arabidopsis and other plants (Silva-Sanchez et al., 2015; Millar et al., 2019) (and in other eukaryotes). Phosphorylation occurs at serine, threonine and tyrosine residues and the distribution of pS, pT and pY is ∼80-85%, ∼10-15%, and 0-5%, respectively in large scale plant (meta) studies (van Wijk et al., 2014; Mergner et al., 2020).

We focused our efforts and resources to build a new PeptideAtlas tool to provide detailed and comprehensive information about protein phosphorylation and phospho-site (p-site) determination. The PeptideAtlas build provides an in-depth view of observed p-sites including statistical significance and associated spectra. All localization p-site probabilities are computed with the TPP tool PTMProphet (Shteynberg et al., 2019) after running iProphet for each dataset. PTMProphet considers all possible permutations of positions of the phosphates reported by Comet and computes Bayesian probabilities that a phosphate is located at each potential STY site based on the subtle differences in spectrum peaks expected for the different permutations. A high probability (*e.g.* P>0.95) indicates a high likelihood that a phosphate was present at a site based on the spectral evidence; a low probability near 0 (e.g. P<0.05) indicates high confidence that a phosphate was not at a site; a probability near 0.5 indicates inability to localize the position of the phosphate with confidence based on the available mass spectrum peaks. Considering only p-sites with a score of P>0.95 or P>0.99 and considering only canonical proteins, the current PeptideAtlas build identified 31988 p-sites for 7778 canonical proteins (44% of the canonical proteins) (Supplemental Dataset 4). The site distribution of S, T and Y was 85% pS, 14% pT and 0.9% pY which is consistent with prior meta-data analysis (van Wijk et al., 2014). When considering only p-sites with 3 or more identifications at P>0.95 or P>0.99, the number of p-sites was 18789 corresponding to 5984 (34%) canonical p-proteins. When considering only p-sites with 3 or more identifications at P>0.99 the number of p-sites was 16028 corresponding to 5565 (31%) canonical proteins. This increased stringency decreased pY to 0.003-0.006% but did not significantly affect the pS and pT ratio. Finally, PeptideAtlas also provides information about the number of phosphorylations per peptide (Supplemental DataSet 4).

Figure 13 demonstrates the PeptideAtlas functionality for p-site analysis by showing examples for the chloroplast Calcium Sensor Protein (CAS) (Figure 13). This is one of many examples in which a protein was identified as a canonical protein and for which phosphorylation was previously demonstrated to have important functional and physiological significance. CAS (AT5G23060.1) was identified with 81% protein sequence coverage and PeptideAtlas shows only one identified p-peptide, *i.e*. the C-terminal peptide SGTKFLPSSD identified 114 times (Figure 13A and 13B). In 84 cases, the p-site at position T380 was identified with the highest significance (0.99 ≤ P ≤ 1.0; in green) and additional observations for this site at lower probabilities (Figure 13C and 13D). There are three serines (S378, S385, S386) in this peptide (no tyrosines), but p-sites were assigned only very low probabilities (see Figure 13C - inset and 13D), indicating high confidence that the detected phosphate was not positioned at those sites. Panel 13D provides a numerical summary of the p-site observations and also provides information about specific sample enrichment for p-peptides based on meta-data information collected from the individual PXD submissions. This shows that the peptide SGTKFLPSSD was 13 times observed in samples that were not enriched for p-peptides and 114 times as a single phosphorylated peptide from enriched samples. Finally, figure 13E shows a detailed peptide view of the phosphorylated peptide SGTKLPSSD and p-site identification scores. The lower panel shows information at individual spectral level with hyperlinked access to the annotated spectra. Together this strongly suggests that CAS is phosphorylated at T380 and very unlikely at S378 (although some spectra are ambiguous). A recent paper suggested T376, S378, and T380 as the major p-sites based on phosphoproteomics and *in vitro* phosphorylation assays on CAS variants (Cutolo et al., 2019), with an earlier study suggesting T380 as the main p-site (Vainonen et al., 2008). PeptideAtlas showed no sequence coverage for T376 despite the very high sequence coverage of the protein (possible peptides with one missed cleavage are SFGT_376_RSGTK or IIPAASRSFGT_376_R or SFGT_376_RSGTK but these were not observed) (Figure 13B,C). All p-peptides that covered S378 indicated either high confidence not at that site, or ambiguous evidence.

**Figure 13.**
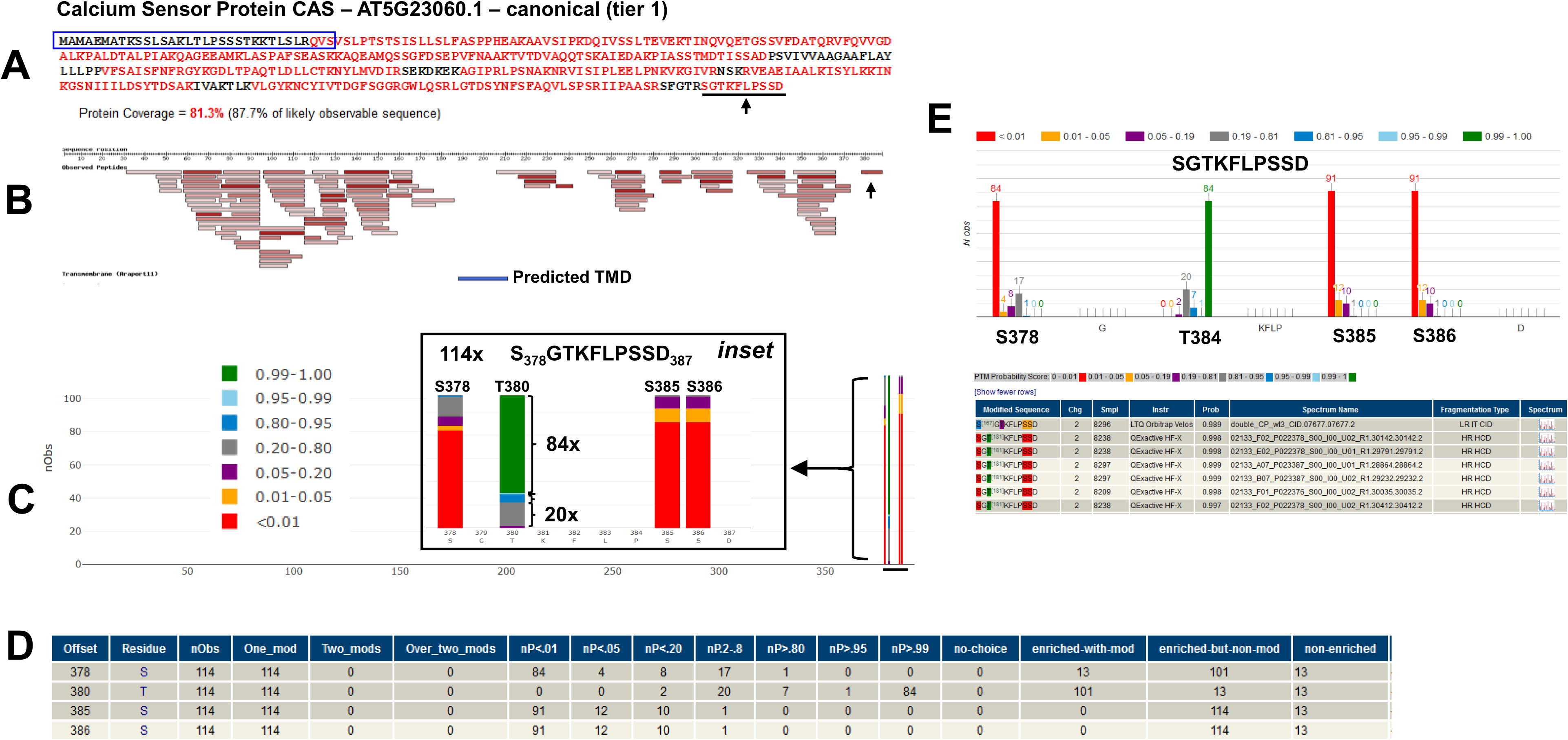
Illustration of the functionality of PeptideAtlas for the determination of phospho-sites based on the example of the thylakoid Calcium Sensor Protein (CAS) AT5G23060.1. **(A)** Coverage by MSMS for the primary sequence (81.3%) with identified residues in red. The predicted cleavable chloroplast signal peptide is indicated in blue, but the spectral evidence suggests the real signal peptide is shorter. **(B)** Matched peptides projected on the primary sequence. Darker red rectangles indicate higher numbers of PSMs for each peptide. The predicted thylakoid transmembrane domain is indicated as a blue rectangle (we note that the N- and C-termini as facing the chloroplast stromal site based on (Cutolo et al., 2019)). **(C)** Probabilities of localization of phosphates on potential sites (as indicated by the colored bars) along the complete protein sequence. The inset provides a close-up of the C-terminal peptide SGTKLPSSD and the frequency of specific p-sites, color-coded by localization probability. This shows *e.g.* that the phosphorylated peptide was observed 114 times and that pT380 was observed 84 times at the highest significance level. **(D)** This small table provides a numerical summary of the p-site observations and information about specific sample enrichment for phospho-peptides based on meta-data information collected from the individual PXD submissions. This shows that the peptide SGTKFLPSSD was 13 times observed in from samples that were not enriched for p-peptides and 114 times as a single p-peptide from enriched samples. Explanation for columns: Offset - residue number from start; Residue - amino acid; nObs - Total observed PTM spectra for the site; One_mod - The number of PSMs with a single phosphate covering the site. Two_mods - The number of PSMs that have two observed phosphates (i.e. doubly phosphorylated). Over_two_mods - The number of PSMs covering the site that have more than two phosphates. nP<.01 - PTMProphet probability < 0.01; nP<.05 - PTMProphet probability >= 0.01 and < 0.05; nP<.20 - PTMProphet probability >= 0.05 and <= 0.20; nP.2-.8 - PTMProphet probability > 0.20 and < 0.80; nP>.80 - PTMProphet probability >= 0.80 and < 0.95; nP>.95 - PTMProphet probability >= 0.95 and < 0.99; nP>.99 - PTMProphet probability >= 0.99; no-choice - Number of PSMs covering this site for which there was no choice in the localization of the PTM. Only one residue was an S, T, or Y; enriched-with-mod - Number of PSMs covering this site with phospho modification on this site, and originating from a phospho-enriched sample; enriched-but-non-mod - Number of PSMs covering this site with no phospho modification anywhere on the peptide, but yet originating from a phospho-enriched sample; non-enriched - Number of PSMs covering this site from a non-enriched sample (phospho not considered in the search). **(E)** Detailed view of the phosphorylated peptide SGTKLPSSD and p-site localization probability distributions. The lower panel shows information at an individual spectral level with hyperlinked access to the annotated spectra.

At a later stage, we will build similar tools for other PTMs, in particular for N-terminal and lysine acetylation, both important physiological PTMs that affect protein localization, protein stability and protein-protein interactions and with functional connections to the metabolic state of the cell through intracellular concentrations of acetyl-coA. In addition to these *in vivo* PTMs, several PTMs are often generated during sample preparation due to exposure to organic solvents (*e.g.* formic acid leading to formylation of Ser, Thr and N-termini), (thio)urea (N-terminal or Lys carbamylation), reducing agents and oxygen, unpolymerized acrylamide (Cys propionamide) and low or high pH (cyclization of N-terminal Gln or Glu into pyro-Glu), reviewed in (Friso and van Wijk, 2015). A large-scale proteomics study on Arabidopsis leaf extracts did address the frequency of PTMs that do not require specific affinity enrichment based on a Dataset of 1.5 million MSMS spectra acquired at 100,000 resolution on an LTQ-Orbitrap instrument followed by error tolerant searches and systematic validation by LC retention time (Zybailov et al., 2009). This showed for example that modification of Met and N-terminal Gln into respectively Met-ox and pyro-Glu showed by far the highest modification frequencies in seedlings (80% of all M observed and 46% of all N-terminal Q), followed by N-terminal formylation (1.5% of all N-termini) most likely induced during sample analysis, as well as deamidation of Asn/Gln (∼1.2% of all observed Asn/Gln). Several of these non-enzymatic PTMs (in particular deamidation, oxidation and formylation) can also occur *in vivo* and therefore cannot be simply dismissed as artifacts but need to be considered as potential regulators. To improve protein coverage and match more MSMS spectra to modified peptides, several PTMs of these were included in all our MS searches (see METHODS).

### Integration with community resources and use of the Arabidopsis Peptide*Atlas*

This first build is freely available in an interactive manner at the PeptideAtlas web site (http://peptideatlas.org/builds/arabidopsis). The results are also made available via web services, allowing easy access to formatted data via external software. We also provide download access to the entire build, which allows anyone to integrate large amounts of the data into their analyses or resource. The build is available as a set of text files, a fully structured XML file, and a MySQL dump that enables easy ingestion into a local MySQL relational database for querying by expert users. Data are already being pulled via web services by UniProtKB and links are established at the protein identifier level (for Araport11 sequences) with TAIR https://www.arabidopsis.org/ and the PPDB http://ppdb.tc.cornell.edu/. The matched peptide data for Araport11 genes in PeptideAtlas are also integrated on a specific track in the Arabidopsis JB browser at https://jbrowse.arabidopsis.org/. We will work with TAIR to further validate and incorporate peptide matched to non-Araport11 sequences, such as those for the sORFs (personal communication Tanya Berardini).

### The next Arabidopsis PeptideAtlas build

The objectives for the next build will be to discover proteins that have so far not been confidently identified in the current build. As was illustrated in figure 11, reasons for the lack of protein identification can include unfavorable physicochemical properties (small, hydrophobic, acidic pI), generally very low copy numbers (*e.g.* for ion channels, and some transcription factors), or if expression is limited to specialized cell types or subcellular locations only present in smaller numbers or under very specific developmental or environmental conditions. We envision three strategies to increase the detection of such proteins, namely i) include PXDs of very specific cell types or specialized subcellular fractions, ii) include PXDs that concern specific protein complexes or protein affinity enrichments, iii) include PXDs that are enriched for specific post-translational modifications. We will also include PXDs that appear to have very high dynamic resolution and sensitivity *e.g.* by using the latest technologies in mass spectrometry and/or sample fractionation.

## Supplemental Data

**Supplemental Text.**

**Supplemental Dataset 1.** Detailed information of the 52 selected PXD files for this first PeptideAtlas build.

**Supplemental Dataset 2.** Identified proteins and unobserved proteins in the PeptideAtlas build and their assignment to the 10-tier system.

**Supplemental Dataset 3.** Evidence for protein identifiers not found in Araport11.

**Supplemental Dataset 4.** Phosphorylation observations in PeptideAtlas.

## ACKNOWLEGDEMENTS

This project is supported by a grant from the National Science Foundation #1922871 to KJVW, EWD and QS. We thank members of the Scientific Advisory board Tanya Berardini, Chris Town, Nicholas Provart, Sixue Chen and Joshua Heazlewood for advice and feedback. We thank Eve Wurtele for sending us her candidate orphan sequences for inclusion in this build.

## AUTHOR CONTRIBUTIONS

TL carried out all MS searches and created the PeptideAtlas build. QS and SSB carried out the physicochemical and functional property analysis of the proteome, supported analysis of sORFs and other non-protein coding identifiers. ZS developed PeptideAtlas interface enhancements and assisted with the PeptideAtlas build process. LM developed PeptideAtlas interface enhancements and developed with dataset annotation tool. ED supervised the PeptideAtlas building process, KJVW contributed to selection of PXDs and all aspects of specific plant biology related issues. EWD and KJVW developed this project, raised the funding, and wrote the paper.

## Parsed Citations

Akter, S., Huang, J., Waszczak, C., Jacques, S., Gevaert, K., Van Breusegem, F., and Messens, J. (2015). Cysteines under ROS attack in plants: a proteomics view. J. Exp. Bot. 66, 2935–2944.

Al-Mohanna, T., Ahsan, N., Bokros, N.T., Dimlioglu, G., Reddy, K.R., Shankle, M., Popescu, G.V., and Popescu, S.C. (2019). Proteomics and Proteogenomics Analysis of Sweetpotato (Ipomoea batatas) Leaf and Root. J Proteome Res 18, 2719–2734.

Ankney, J.A., Muneer, A., and Chen, X. (2018). Relative and Absolute Quantitation in Mass Spectrometry-Based Proteomics. Annu Rev Anal Chem(Palo Alto Calif) 11, 49–77.

Augustine, R.C., and Vierstra, R.D. (2018). SUMOylation: re-wiring the plant nucleus during stress and development. Curr. Opin. Plant Biol. 45, 143–154.

Balmant, K.M., Zhang, T., and Chen, S. (2016). Protein Phosphorylation and Redox Modification in Stomatal Guard Cells. Front Physiol 7, 26.

Bislev, S.L., Deutsch, E.W., Sun, Z., Farrah, T., Aebersold, R., Moritz, R.L., Bendixen, E., and Codrea, M.C. (2012). ABovine PeptideAtlas of milk and mammary gland proteomes. Proteomics 12, 2895–2899.

Blencowe, B.J. (2017). The Relationship between Alternative Splicing and Proteomic Complexity. Trends Biochem. Sci. 42, 407–408.

Bolter, B., Mitterreiter, M.J., Schwenkert, S., Finkemeier, I., and Kunz, H.H. (2020). The topology of plastid inner envelope potassium cation efflux antiporter KEA1 provides new insights into its regulatory features. Photosynth Res 145, 43–54.

Calderon-Celis, F., Encinar, J.R., and Sanz-Medel, A. (2018). Standardization approaches in absolute quantitative proteomics with mass spectrometry. Mass Spectrom Rev 37, 715–737.

Castellana, N.E., Shen, Z., He, Y., Walley, J.W., Cassidy, C.J., Briggs, S.P., and Bafna, V. (2014). An automated proteogenomic method uses mass spectrometry to reveal novel genes in Zea mays. Mol. Cell. Proteomics 13, 157–167.

Chambers, M.C., Maclean, B., Burke, R., Amodei, D., Ruderman, D.L., Neumann, S., Gatto, L., Fischer, B., Pratt, B., Egertson, J., Hoff, K., Kessner, D., Tasman, N., Shulman, N., Frewen, B., Baker, T.A., Brusniak, M.Y., Paulse, C., Creasy, D., Flashner, L., Kani, K., Moulding, C., Seymour, S.L., Nuwaysir, L.M., Lefebvre, B., Kuhlmann, F., Roark, J., Rainer, P., Detlev, S., Hemenway, T., Huhmer, A., Langridge, J., Connolly, B., Chadick, T., Holly, K., Eckels, J., Deutsch, E.W., Moritz, R.L., Katz, J.E., Agus, D.B., MacCoss, M., Tabb, D.L., and Mallick, P. (2012). Across-platformtoolkit for mass spectrometry and proteomics. Nat. Biotechnol. 30, 918–920.

Chang, W.W., Huang, L., Shen, M., Webster, C., Burlingame, A.L., and Roberts, J.K. (2000). Patterns of protein synthesis and tolerance of anoxia in root tips of maize seedlings acclimated to a low-oxygen environment, and identification of proteins by mass spectrometry. Plant Physiol. 122, 295–318.

Chapman, B., and Bellgard, M. (2017). Plant Proteogenomics: Improvements to the Grapevine Genome Annotation. Proteomics 17.

Chen, C., Liu, X., Zheng, W., Zhang, L., Yao, J., and Yang, P. (2014). Screening of missing proteins in the human liver proteome by improved MRM-approach-based targeted proteomics. J Proteome Res 13, 1969–1978.

Chen, M.X., Zhu, F.Y., Gao, B., Ma, K.L., Zhang, Y., Fernie, A.R., Chen, X., Dai, L., Ye, N.H., Zhang, X., Tian, Y., Zhang, D., Xiao, S., Zhang, J., and Liu, Y.G. (2020). Full-Length Transcript-Based Proteogenomics of Rice Improves Its Genome and Proteome Annotation. Plant Physiol. 182, 1510–1526.

Cheng, C.Y., Krishnakumar, V., Chan, A.P., Thibaud-Nissen, F., Schobel, S., and Town, C.D. (2017). Araport11: a complete reannotation of the Arabidopsis thaliana reference genome. Plant J. 89, 789–804.

Cottingham, K. (2009). Two are not always better than one. J Proteome Res 8, 4172.

Creasy, D.M., and Cottrell, J.S. (2004). Unimod: Protein modifications for mass spectrometry. Proteomics 4, 1534–1536.

Cutolo, E., Parvin, N., Ruge, H., Pirayesh, N., Roustan, V., Weckwerth, W., Teige, M., Grieco, M., Larosa, V., and Vothknecht, U.C. (2019). The High Light Response in Arabidopsis Requires the Calcium Sensor Protein CAS, a Target of STN7- and STN8-Mediated Phosphorylation. Front Plant Sci 10, 974.

Desiere, F., Deutsch, E.W., Nesvizhskii, A.I., Mallick, P., King, N.L., Eng, J.K., Aderem, A., Boyle, R., Brunner, E., Donohoe, S., Fausto, N., Hafen, E., Hood, L., Katze, M.G., Kennedy, K.A., Kregenow, F., Lee, H., Lin, B., Martin, D., Ranish, J.A., Rawlings, D.J., Samelson, L.E., Shiio, Y., Watts, J.D., Wollscheid, B., Wright, M.E., Yan, W., Yang, L., Yi, E.C., Zhang, H., and Aebersold, R. (2005). Integration with the human genome of peptide sequences obtained by high-throughput mass spectrometry. Genome Biol 6, R9.

Deutsch, E.W., Mendoza, L., Shteynberg, D., Slagel, J., Sun, Z., and Moritz, R.L. (2015). Trans-Proteomic Pipeline, a standardized data processing pipeline for large-scale reproducible proteomics informatics. Proteomics Clin Appl 9, 745–754.

Deutsch, E.W., Sun, Z., Campbell, D.S., Binz, P.A., Farrah, T., Shteynberg, D., Mendoza, L., Omenn, G.S., and Moritz, R.L. (2016a). Tiered Human Integrated Sequence Search Databases for Shotgun Proteomics. J Proteome Res 15, 4091–4100.

Deutsch, E.W., Lane, L., Overall, C.M., Bandeira, N., Baker, M.S., Pineau, C., Moritz, R.L., Corrales, F., Orchard, S., Van Eyk, J.E., Paik, Y.K., Weintraub, S.T., Vandenbrouck, Y., and Omenn, G.S. (2019). Human Proteome Project Mass Spectrometry Data Interpretation Guidelines 3.0. J Proteome Res 18, 4108–4116.

Deutsch, E.W., Overall, C.M., Van Eyk, J.E., Baker, M.S., Paik, Y.K., Weintraub, S.T., Lane, L., Martens, L., Vandenbrouck, Y., Kusebauch, U., Hancock, W.S., Hermjakob, H., Aebersold, R., Moritz, R.L., and Omenn, G.S. (2016b). Human Proteome Project Mass Spectrometry Data Interpretation Guidelines 2.1. J Proteome Res 15, 3961–3970.

Deutsch, E.W., Orchard, S., Binz, P.A., Bittremieux, W., Eisenacher, M., Hermjakob, H., Kawano, S., Lam, H., Mayer, G., Menschaert, G., Perez-Riverol, Y., Salek, R.M., Tabb, D.L., Tenzer, S., Vizcaino, J.A., Walzer, M., and Jones, A.R. (2017a). Proteomics Standards Initiative: Fifteen Years of Progress and Future Work. J Proteome Res 16, 4288–4298.

Deutsch, E.W., Csordas, A., Sun, Z., Jarnuczak, A., Perez-Riverol, Y., Ternent, T., Campbell, D.S., Bernal-Llinares, M., Okuda, S., Kawano, S., Moritz, R.L., Carver, J.J., Wang, M., Ishihama, Y., Bandeira, N., Hermjakob, H., and Vizcaino, J.A. (2017b). The ProteomeXchange consortiumin 2017: supporting the cultural change in proteomics public data deposition. Nucleic Acids Res. 45, D1100–D1106.

Deutsch, E.W., Bandeira, N., Sharma, V., Perez-Riverol, Y., Carver, J.J., Kundu, D.J., Garcia-Seisdedos, D., Jarnuczak, A.F., Hewapathirana, S., Pullman, B.S., Wertz, J., Sun, Z., Kawano, S., Okuda, S., Watanabe, Y., Hermjakob, H., MacLean, B., MacCoss, M.J., Zhu, Y., Ishihama, Y., and Vizcaino, J.A. (2020). The ProteomeXchange consortiumin 2020: enabling ‘big data’ approaches in proteomics. Nucleic Acids Res. 48, D1145–D1152.

Eliuk, S., and Makarov, A. (2015). Evolution of Orbitrap Mass Spectrometry Instrumentation. Annu Rev Anal Chem(Palo Alto Calif) 8, 61–80.

Eng, J.K., and Deutsch, E.W. (2020). Extending Comet for Global Amino Acid Variant and Post-Translational Modification Analysis Using the PSI Extended FASTA Format. Proteomics, e1900362.

Farrah, T., Deutsch, E.W., Omenn, G.S., Campbell, D.S., Sun, Z., Bletz, J.A., Mallick, P., Katz, J.E., Malmstrom, J., Ossola, R., Watts, J.D., Lin, B., Zhang, H., Moritz, R.L., and Aebersold, R. (2011). Ahigh-confidence human plasma proteome reference set with estimated concentrations in PeptideAtlas. Mol. Cell. Proteomics 10, M110 006353.

Friso, G., and van Wijk, K.J. (2015). Posttranslational Protein Modifications in Plant Metabolism. Plant Physiol. 169, 1469–1487.

Fuchs, P., Rugen, N., Carrie, C., Elsasser, M., Finkemeier, I., Giese, J., Hildebrandt, T.M., Kuhn, K., Maurino, V.G., Ruberti, C., Schallenberg-Rudinger, M., Steinbeck, J., Braun, H.P., Eubel, H., Meyer, E.H., Muller-Schussele, S.J., and Schwarzlander, M. (2020). Single organelle function and organization as estimated from Arabidopsis mitochondrial proteomics. Plant J. 101, 420–441.

Fussl, M., Lassowskat, I., Nee, G., Koskela, M.M., Brunje, A., Tilak, P., Giese, J., Leister, D., Mulo, P., Schwarzer, D., and Finkemeier, I. (2018). Beyond Histones: New Substrate Proteins of Lysine Deacetylases in Arabidopsis Nuclei. Front Plant Sci 9, 461.

Germain, A., Hanson, M.R., and Bentolila, S. (2015). High-throughput quantification of chloroplast RNAediting extent using multiplex RT-PCR mass spectrometry. Plant J. 83, 546–554.

Giglione, C., and Meinnel, T. (2020). Evolution-Driven Versatility of N Terminal Acetylation in Photoautotrophs. Trends Plant Sci.

Hanada, K., Zhang, X., Borevitz, J.O., Li, W.H., and Shiu, S.H. (2007). Alarge number of novel coding small open reading frames in the intergenic regions of the Arabidopsis thaliana genome are transcribed and/or under purifying selection. Genome Res. 17, 632–640.

Hartl, M., Fussl, M., Boersema, P.J., Jost, J.O., Kramer, K., Bakirbas, A., Sindlinger, J., Plochinger, M., Leister, D., Uhrig, G., Moorhead, G.B., Cox, J., Salvucci, M.E., Schwarzer, D., Mann, M., and Finkemeier, I. (2017). Lysine acetylome profiling uncovers novel histone deacetylase substrate proteins in Arabidopsis. Mol. Syst. Biol. 13, 949.

Hazarika, R.R., De Coninck, B., Yamamoto, L.R., Martin, L.R., Cammue, B.P., and van Noort, V. (2017). ARA-PEPs: a repository of putative sORF-encoded peptides in Arabidopsis thaliana. BMC Bioinformatics 18, 37.

Hesselager, M.O., Codrea, M.C., Sun, Z., Deutsch, E.W., Bennike, T.B., Stensballe, A., Bundgaard, L., Moritz, R.L., and Bendixen, E. (2016). The Pig PeptideAtlas: Aresource for systems biology in animal production and biomedicine. Proteomics 16, 634–644.

Hosp, F., Lassowskat, I., Santoro, V., De Vleesschauwer, D., Fliegner, D., Redestig, H., Mann, M., Christian, S., Hannah, M.A., and Finkemeier, I. (2017). Lysine acetylation in mitochondria: Frominventory to function. Mitochondrion 33, 58–71.

Hsu, P.Y., and Benfey, P.N. (2018). Small but Mighty: Functional Peptides Encoded by Small ORFs in Plants. Proteomics 18, e1700038.

Hsu, P.Y., Calviello, L., Wu, H.L., Li, F.W., Rothfels, C.J., Ohler, U., and Benfey, P.N. (2016). Super-resolution ribosome profiling reveals unannotated translation events in Arabidopsis. Proc Natl Acad Sci U S A 113, E7126–E7135.

Huang, M.D., Chen, T.L., and Huang, A.H. (2013). Abundant type III lipid transfer proteins in Arabidopsis tapetumare secreted to the locule and become a constituent of the pollen exine. Plant Physiol. 163, 1218–1229.

Hulstaert, N., Shofstahl, J., Sachsenberg, T., Walzer, M., Barsnes, H., Martens, L., and Perez-Riverol, Y. (2020). ThermoRawFileParser: Modular, Scalable, and Cross-Platform RAW File Conversion. J Proteome Res 19, 537–542.

Initiative, T.A.G. (2000). Analysis of the genome sequence of the flowering plant Arabidopsis thaliana. Nature 408, 796–815.

Kage, U., Powell, J.J., Gardiner, D.M., and Kazan, K. (2020). Ribosome Profiling in Plants: What is NOT lost in translation? J. Exp. Bot.

Kaur, J., Sebastian, J., and Siddiqi, I. (2006). The Arabidopsis-mei2-like genes play a role in meiosis and vegetative growth in Arabidopsis. Plant Cell 18, 545–559.

Keller, A., Nesvizhskii, A.I., Kolker, E., and Aebersold, R. (2002). Empirical statistical model to estimate the accuracy of peptide identifications made by MS/MS and database search. Anal Chem 74, 5383–5392.

Keller, A., Eng, J., Zhang, N., Li, X.J., and Aebersold, R. (2005). Auniformproteomics MS/MS analysis platformutilizing open XML file formats. Mol. Syst. Biol. 1, 2005 0017.

King, N.L., Deutsch, E.W., Ranish, J.A., Nesvizhskii, A.I., Eddes, J.S., Mallick, P., Eng, J., Desiere, F., Flory, M., Martin, D.B., Kim, B., Lee, H., Raught, B., and Aebersold, R. (2006). Analysis of the Saccharomyces cerevisiae proteome with PeptideAtlas. Genome Biol 7, R106.

Kiraga, J., Mackiewicz, P., Mackiewicz, D., Kowalczuk, M., Biecek, P., Polak, N., Smolarczyk, K., Dudek, M.R., and Cebrat, S. (2007). The relationships between the isoelectric point and: length of proteins, taxonomy and ecology of organisms. BMC Genomics 8, 163.

Lamesch, P., Berardini, T.Z., Li, D., Swarbreck, D., Wilks, C., Sasidharan, R., Muller, R., Dreher, K., Alexander, D.L., Garcia-Hernandez, M., Karthikeyan, A.S., Lee, C.H., Nelson, W.D., Ploetz, L., Singh, S., Wensel, A., and Huala, E. (2012). The Arabidopsis Information Resource (TAIR): improved gene annotation and new tools. Nucleic Acids Res. 40, D1202–1210.

Li, W., O’Neill, K.R., Haft, D.H., DiCuccio, M., Chetvernin, V., Badretdin, A., Coulouris, G., Chitsaz, F., Derbyshire, M.K., Durkin, A.S., Gonzales, N.R., Gwadz, M., Lanczycki, C.J., Song, J.S., Thanki, N., Wang, J., Yamashita, R.A., Yang, M., Zheng, C., Marchler-Bauer, A., and Thibaud-Nissen, F. (2020). RefSeq: expanding the Prokaryotic Genome Annotation Pipeline reach with protein family model curation. Nucleic Acids Res.

Linster, E., and Wirtz, M. (2018). N-terminal acetylation: an essential protein modification emerges as an important regulator of stress responses. J. Exp. Bot. 69, 4555–4568.

Makarov, A. (2019). Orbitrap journey: taming the ion rings. Nat Commun 10, 3743.

Marino, G., Eckhard, U., and Overall, C.M. (2015). Protein Termini and Their Modifications Revealed by Positional Proteomics. ACS Chem Biol 10, 1754–1764.

Martens, L., Chambers, M., Sturm, M., Kessner, D., Levander, F., Shofstahl, J., Tang, W.H., Rompp, A., Neumann, S., Pizarro, A.D., Montecchi-Palazzi, L., Tasman, N., Coleman, M., Reisinger, F., Souda, P., Hermjakob, H., Binz, P.A., and Deutsch, E.W. (2011). mzML--a community standard for mass spectrometry data. Mol. Cell. Proteomics 10, R110 000133.

Mayer, G., Montecchi-Palazzi, L., Ovelleiro, D., Jones, A.R., Binz, P.A., Deutsch, E.W., Chambers, M., Kallhardt, M., Levander, F., Shofstahl, J., Orchard, S., Vizcaino, J.A., Hermjakob, H., Stephan, C., Meyer, H.E., Eisenacher, M., and Group, H.-P. (2013). The HUPO proteomics standards initiative-mass spectrometry controlled vocabulary. Database (Oxford) 2013, bat009.

Mayfield, J.A., Fiebig, A., Johnstone, S.E., and Preuss, D. (2001). Gene families fromthe Arabidopsis thaliana pollen coat proteome. Science 292, 2482–2485.

McCord, J., Sun, Z., Deutsch, E.W., Moritz, R.L., and Muddiman, D.C. (2017). The PeptideAtlas of the Domestic Laying Hen. J Proteome Res 16, 1352–1363.

Mergner, J., Frejno, M., List, M., Papacek, M., Chen, X., Chaudhary, A., Samaras, P., Richter, S., Shikata, H., Messerer, M., Lang, D., Altmann, S., Cyprys, P., Zolg, D.P., Mathieson, T., Bantscheff, M., Hazarika, R.R., Schmidt, T., Dawid, C., Dunkel, A., Hofmann, T., Sprunck, S., Falter-Braun, P., Johannes, F., Mayer, K.F.X., Jurgens, G., Wilhelm, M., Baumbach, J., Grill, E., Schneitz, K., Schwechheimer, C., and Kuster, B. (2020). Mass-spectrometry-based draft of the Arabidopsis proteome. Nature 579, 409–414.

Millar, A.H., Sweetlove, L.J., Giege, P., and Leaver, C.J. (2001). Analysis of the Arabidopsis mitochondrial proteome. Plant Physiol. 127, 1711–1727.

Millar, A.H., Heazlewood, J.L., Giglione, C., Holdsworth, M.J., Bachmair, A., and Schulze, W.X. (2019). The Scope, Functions, and Dynamics of Posttranslational Protein Modifications. Annu. Rev. Plant Biol. prepublication online, 119–151.

Misra, B.B. (2018). Updates on resources, software tools, and databases for plant proteomics in 2016-2017. Electrophoresis 39, 1543–1557.

Miura, K., and Hasegawa, P.M. (2010). Sumoylation and other ubiquitin-like post-translational modifications in plants. Trends Cell Biol. 20, 223–232.

Moller, I.M., Igamberdiev, A.U., Bykova, N.V., Finkemeier, I., Rasmusson, A.G., and Schwarzlander, M. (2020). Matrix Redox Physiology Governs the Regulation of Plant Mitochondrial Metabolismthrough Posttranslational Protein Modifications. Plant Cell 32, 573–594.

Montecchi-Palazzi, L., Beavis, R., Binz, P.A., Chalkley, R.J., Cottrell, J., Creasy, D., Shofstahl, J., Seymour, S.L., and Garavelli, J.S. (2008). The PSI-MOD community standard for representation of protein modification data. Nat. Biotechnol. 26, 864–866.

Omenn, G.S., Lane, L., Overall, C.M., Corrales, F.J., Schwenk, J.M., Paik, Y.K., Van Eyk, J.E., Liu, S., Pennington, S., Snyder, M.P., Baker, M.S., and Deutsch, E.W. (2019). Progress on Identifying and Characterizing the Human Proteome: 2019 Metrics fromthe HUPO Human Proteome Project. J Proteome Res 18, 4098–4107.

Omenn, G.S., Lane, L., Overall, C.M., Cristea, I.M., Corrales, F.J., Lindskog, C., Paik, Y.K., Van Eyk, J.E., Liu, S., Pennington, S.R., Snyder, M.P., Baker, M.S., Bandeira, N., Aebersold, R., Moritz, R.L., and Deutsch, E.W. (2020). Research on the Human Proteome Reaches a Major Milestone: >90% of Predicted Human Proteins Now Credibly Detected, According to the HUPO Human Proteome Project. J Proteome Res 19, 4735–4746.

Orchard, S., Hermjakob, H., and Apweiler, R. (2003). The proteomics standards initiative. Proteomics 3, 1374–1376.

Peltier, J.B., Ytterberg, J., Liberles, D.A., Roepstorff, P., and van Wijk, K.J. (2001). Identification of a 350-kDa ClpP protease complex with 10 different Clp isoforms in chloroplasts of Arabidopsis thaliana. J. Biol. Chem. 276, 16318–16327.

Peltier, J.B., Friso, G., Kalume, D.E., Roepstorff, P., Nilsson, F., Adamska, I., and van Wijk, K.J. (2000). Proteomics of the Chloroplast. Systematic identification and targeting analysis of lumenal and peripheral thylakoid proteins. Plant Cell 12, 319–342.

Peltier, J.B., Emanuelsson, O., Kalume, D.E., Ytterberg, J., Friso, G., Rudella, A., Liberles, D.A., Soderberg, L., Roepstorff, P., von Heijne, G., and van Wijk, K.J. (2002). Central Functions of the Lumenal and Peripheral Thylakoid Proteome of Arabidopsis Determined by Experimentation and Genome-Wide Prediction. Plant Cell 14, 211–236.

Perez-Riverol, Y., Csordas, A., Bai, J., Bernal-Llinares, M., Hewapathirana, S., Kundu, D.J., Inuganti, A., Griss, J., Mayer, G., Eisenacher, M., Perez, E., Uszkoreit, J., Pfeuffer, J., Sachsenberg, T., Yilmaz, S., Tiwary, S., Cox, J., Audain, E., Walzer, M., Jarnuczak, A.F., Ternent, T., Brazma, A., and Vizcaino, J.A. (2018). The PRIDE database and related tools and resources in 2019: improving support for quantification data. Nucleic Acids Res.

Provart, N.J., Alonso, J., Assmann, S.M., Bergmann, D., Brady, S.M., Brkljacic, J., Browse, J., Chapple, C., Colot, V., Cutler, S., Dangl, J., Ehrhardt, D., Friesner, J.D., Frommer, W.B., Grotewold, E., Meyerowitz, E., Nemhauser, J., Nordborg, M., Pikaard, C., Shanklin, J., Somerville, C., Stitt, M., Torii, K.U., Waese, J., Wagner, D., and McCourt, P. (2016). 50 years of Arabidopsis research: highlights and future directions. New Phytol. 209, 921–944.

Qi, Y., Wang, X., Lei, P., Li, H., Yan, L., Zhao, J., Meng, J., Shao, J., An, L., Yu, F., and Liu, X. (2020). The chloroplast metalloproteases VAR2 and EGY1 act synergistically to regulate chloroplast development in Arabidopsis. J. Biol. Chem. 295, 1036–1046.

Quesneville, H. (2020). Twenty years of transposable element analysis in the Arabidopsis thaliana genome. Mob DNA 11, 28.

Ren, Z., Qi, D., Pugh, N., Li, K., Wen, B., Zhou, R., Xu, S., Liu, S., and Jones, A.R. (2019). Improvements to the Rice Genome Annotation Through Large-Scale Analysis of RNA-Seq and Proteomics Data Sets. Mol. Cell. Proteomics 18, 86–98.

Roberts, I., Smith, S., De Rybel, B., Van Den Broeke, J., Smet, W., De Cokere, S., Mispelaere, M., De Smet, I., and Beeckman, T. (2013). The CEP family in land plants: evolutionary analyses, expression studies, and role in Arabidopsis shoot development. J. Exp. Bot. 64, 5371–5381.

Rowland, E., Kim, J., Bhuiyan, N.H., and van Wijk, K.J. (2015). The Arabidopsis Chloroplast Stromal N-Terminome: Complexities of Amino-Terminal Protein Maturation and Stability. Plant Physiol. 169, 1881–1896.

Ruiz-May, E., Segura-Cabrera, A., Elizalde-Contreras, J.M., Shannon, L.M., and Loyola-Vargas, V.M. (2019). Arecent advance in the intracellular and extracellular redox post-translational modification of proteins in plants. J Mol Recognit 32, e2754.

Salvi, D., Bournais, S., Moyet, L., Bouchnak, I., Kuntz, M., Bruley, C., and Rolland, N. (2018). AT_CHLORO: The First Step When Looking for Information About Subplastidial Localization of Proteins. Methods Mol Biol 1829, 395–406.

San Clemente, H., and Jamet, E. (2015). WallProtDB, a database resource for plant cell wall proteomics. Plant Methods 11, 2.

Sandalio, L.M., Gotor, C., Romero, L.C., and Romero-Puertas, M.C. (2019). Multilevel Regulation of Peroxisomal Proteome by Post-Translational Modifications. Int J Mol Sci 20.

Sato, S., Nakamura, Y., Kaneko, T., Asamizu, E., and Tabata, S. (1999). Complete structure of the chloroplast genome of Arabidopsis thaliana. DNARes. 6, 283–290.

Schubert, M., Petersson, U.A., Haas, B.J., Funk, C., Schröder, W.P., and Kieselbach, T. (2002). Proteome map of the chloroplast lumen of Arabidopsis thaliana. J. Biol. Chem. 277, 8354–8365.

Schulze, W.X., Yao, Q., and Xu, D. (2015). Databases for plant phosphoproteomics. Methods Mol Biol 1306, 207–216.

Schwartz, R., Ting, C.S., and King, J. (2001). Whole proteome pI values correlate with subcellular localizations of proteins for organisms within the three domains of life. Genome Res. 11, 703–709.

Shteynberg, D., Deutsch, E.W., Lam, H., Eng, J.K., Sun, Z., Tasman, N., Mendoza, L., Moritz, R.L., Aebersold, R., and Nesvizhskii, A.I. (2011). iProphet: multi-level integrative analysis of shotgun proteomic data improves peptide and protein identification rates and error estimates. Mol. Cell. Proteomics 10, M111 007690.

Shteynberg, D.D., Deutsch, E.W., Campbell, D.S., Hoopmann, M.R., Kusebauch, U., Lee, D., Mendoza, L., Midha, M.K., Sun, Z., Whetton, A.D., and Moritz, R.L. (2019). PTMProphet: Fast and Accurate Mass Modification Localization for the Trans-Proteomic Pipeline. J Proteome Res 18, 4262–4272.

Silva-Sanchez, C., Li, H., and Chen, S. (2015). Recent advances and challenges in plant phosphoproteomics. Proteomics 15, 1127–1141.

Slagel, J., Mendoza, L., Shteynberg, D., Deutsch, E.W., and Moritz, R.L. (2015). Processing shotgun proteomics data on the Amazon cloud with the trans-proteomic pipeline. Mol. Cell. Proteomics 14, 399–404.

Sloan, D.B., Wu, Z., and Sharbrough, J. (2018). Correction of Persistent Errors in Arabidopsis Reference Mitochondrial Genomes. Plant Cell 30, 525–527.

Small, I.D., Schallenberg-Rudinger, M., Takenaka, M., Mireau, H., and Ostersetzer-Biran, O. (2020). Plant organellar RNAediting: what 30 years of research has revealed. Plant J. 101, 1040–1056.

Staes, A., Impens, F., Van Damme, P., Ruttens, B., Goethals, M., Demol, H., Timmerman, E., Vandekerckhove, J., and Gevaert, K. (2011). Selecting protein N-terminal peptides by combined fractional diagonal chromatography. Nat Protoc 6, 1130–1141.

Stecker, K.E., Minkoff, B.B., and Sussman, M.R. (2014). Phosphoproteomic Analyses Reveal Early Signaling Events in the Osmotic Stress Response. Plant Physiol. 165, 1171–1187.

Sun, Q., Zybailov, B., Majeran, W., Friso, G., Olinares, P.D., and van Wijk, K.J. (2009). PPDB, the Plant Proteomics Database at Cornell. Nucleic Acids Res. 37, D969–974.

Takahashi, F., Hanada, K., Kondo, T., and Shinozaki, K. (2019). Hormone-like peptides and small coding genes in plant stress signaling and development. Curr. Opin. Plant Biol. 51, 88–95.

Takahashi, H., Hayashi, N., Hiragori, Y., Sasaki, S., Motomura, T., Yamashita, Y., Naito, S., Takahashi, A., Fuse, K., Satou, K., Endo, T., Kojima, S., and Onouchi, H. (2020). Comprehensive genome-wide identification of angiospermupstream ORFs with peptide sequences conserved in various taxonomic ranges using a novel pipeline, ESUCA. BMC Genomics 21, 260.

Takenaka, M., Zehrmann, A., Verbitskiy, D., Hartel, B., and Brennicke, A. (2013). RNAediting in plants and its evolution. Annu. Rev. Genet. 47, 335–352.

Tan, B.C., Lim, Y.S., and Lau, S.E. (2017). Proteomics in commercial crops: An overview. J Proteomics 169, 176–188.

Tanz, S.K., Castleden, I., Hooper, C.M., Vacher, M., Small, I., and Millar, H.A. (2013). SUBA3: a database for integrating experimentation and prediction to define the SUBcellular location of proteins in Arabidopsis. Nucleic Acids Res. 41, D1185–1191.

Tress, M.L., Abascal, F., and Valencia, A. (2017). Alternative Splicing May Not Be the Key to Proteome Complexity. Trends Biochem. Sci. 42, 98–110.

UniProt, C. (2020). UniProt: the universal protein knowledgebase in 2021. Nucleic Acids Res.

Vainonen, J.P., Sakuragi, Y., Stael, S., Tikkanen, M., Allahverdiyeva, Y., Paakkarinen, V., Aro, E., Suorsa, M., Scheller, H.V., Vener, A.V., and Aro, E.M. (2008). Light regulation of CaS, a novel phosphoprotein in the thylakoid membrane of Arabidopsis thaliana. FEBS J. 275, 1767–1777.

van Wijk, K.J. (2000). Proteomics of the chloroplast: experimentation and prediction. Trends Plant Sci. 5, 420–425.

van Wijk, K.J., Friso, G., Walther, D., and Schulze, W.X. (2014). Meta-Analysis of Arabidopsis thaliana Phospho-Proteomics Data Reveals Compartmentalization of Phosphorylation Motifs. Plant Cell 26, 2367–2389.

Vandenbrouck, Y., Lane, L., Carapito, C., Duek, P., Rondel, K., Bruley, C., Macron, C., Gonzalez de Peredo, A., Coute, Y., Chaoui, K., Com, E., Gateau, A., Hesse, A.M., Marcellin, M., Mear, L., Mouton-Barbosa, E., Robin, T., Burlet-Schiltz, O., Cianferani, S., Ferro, M., Freour, T., Lindskog, C., Garin, J., and Pineau, C. (2016). Looking for Missing Proteins in the Proteome of Human Spermatozoa: An Update. J Proteome Res 15, 3998–4019.

Vanderschuren, H., Lentz, E., Zainuddin, I., and Gruissem, W. (2013). Proteomics of model and crop plant species: status, current limitations and strategic advances for crop improvement. J Proteomics 93, 5–19.

Vialas, V., Sun, Z., Loureiro y Penha, C.V., Carrascal, M., Abian, J., Monteoliva, L., Deutsch, E.W., Aebersold, R., Moritz, R.L., and Gil, C. (2014). ACandida albicans PeptideAtlas. J Proteomics 97, 62–68.

Vierstra, R.D. (2012). The expanding universe of ubiquitin and ubiquitin-like modifiers. Plant Physiol. 160, 2–14.

Vizcaino, J.A., Deutsch, E.W., Wang, R., Csordas, A., Reisinger, F., Rios, D., Dianes, J.A., Sun, Z., Farrah, T., Bandeira, N., Binz, P.A., Xenarios, I., Eisenacher, M., Mayer, G., Gatto, L., Campos, A., Chalkley, R.J., Kraus, H.J., Albar, J.P., Martinez-Bartolome, S., Apweiler, R., Omenn, G.S., Martens, L., Jones, A.R., and Hermjakob, H. (2014). ProteomeXchange provides globally coordinated proteomics data submission and dissemination. Nat. Biotechnol. 32, 223–226.

Vu, L.D., Gevaert, K., and De Smet, I. (2018). Protein Language: Post-Translational Modifications Talking to Each Other. Trends Plant Sci. 23, 1068–1080.

Walley, J.W., and Briggs, S.P. (2015). Dual use of peptide mass spectra: Protein atlas and genome annotation. Curr Plant Biol 2, 21–24.

Wang, S., Tian, L., Liu, H., Li, X., Zhang, J., Chen, X., Jia, X., Zheng, X., Wu, S., Chen, Y., Yan, J., and Wu, L. (2020). Large-Scale Discovery of Non-conventional Peptides in Maize and Arabidopsis through an Integrated Peptidogenomic Pipeline. Mol Plant 13, 1078–1093.

Waszczak, C., Akter, S., Jacques, S., Huang, J., Messens, J., and Van Breusegem, F. (2015). Oxidative post-translational modifications of cysteine residues in plant signal transduction. J. Exp. Bot. 66, 2923–2934.

Willems, P., Horne, A., Van Parys, T., Goormachtig, S., De Smet, I., Botzki, A., Van Breusegem, F., and Gevaert, K. (2019). The Plant PTM Viewer, a central resource for exploring plant protein modifications. Plant J. 99, 752–762.

Willems, P., Ndah, E., Jonckheere, V., Stael, S., Sticker, A., Martens, L., Van Breusegem, F., Gevaert, K., and Van Damme, P. (2017). N-terminal Proteomics Assisted Profiling of the Unexplored Translation Initiation Landscape in Arabidopsis thaliana. Mol. Cell. Proteomics 16, 1064–1080.

Ytterberg, J., Peltier, J.B., Friso, G. and van Wijk. K.J.. (2002). Identification and Analysis of the Thylakoid Membrane Proteome of Arabidopsis thaliana by Sequential Organic Solvent Extraction, Gel Based Protein Separation, RP-HPLC, MALDI-TOF MS and CapLC-Q-TOF MS. In American Society for Mass Spectrometry (Orlando, Florida).

Zhang, H., Liu, P., Guo, T., Zhao, H., Bensaddek, D., Aebersold, R., and Xiong, L. (2019). Arabidopsis proteome and the mass spectral assay library. Sci Data 6, 278.

Zhu, F.Y., Chen, M.X., Ye, N.H., Shi, L., Ma, K.L., Yang, J.F., Cao, Y.Y., Zhang, Y., Yoshida, T., Fernie, A.R., Fan, G.Y., Wen, B., Zhou, R., Liu, T.Y., Fan, T., Gao, B., Zhang, D., Hao, G.F., Xiao, S., Liu, Y.G., and Zhang, J. (2017). Proteogenomic analysis reveals alternative splicing and translation as part of the abscisic acid response in Arabidopsis seedlings. Plant J. 91, 518–533.

Zybailov, B., Rutschow, H., Friso, G., Rudella, A., Emanuelsson, O., Sun, Q., and van Wijk, K.J. (2008). Sorting signals, N-terminal modifications and abundance of the chloroplast proteome. PLoS ONE 3, e1994.

